# A HYPOMETABOLIC DEFENSE STRATEGY AGAINST *PLASMODIUM* INFECTION

**DOI:** 10.1101/2021.09.08.459402

**Authors:** Susana Ramos, Temitope W. Ademolue, Elisa Jentho, Qian Wu, Joel Guerra, Rui Martins, Gil Pires, Sebastian Weis, Ana Rita Carlos, Inês Mahú, Elsa Seixas, Denise Duarte, Fabienne Rajas, Sílvia Cardoso, António G. G. Sousa, Jingtao Lilue, Gilles Mithieux, Fátima Nogueira, Miguel P. Soares

## Abstract

Hypoglycemia is a clinical hallmark of severe malaria, the often-lethal presentation of *Plasmodium falciparum* infection of humans. Here we report that mice reduce blood glucose levels in response to *Plasmodium* infection via a coordinated response whereby labile heme, an alarmin produced via hemolysis, induces anorexia and represses hepatic glucose production (HGP). While protective against unfettered immune-mediated inflammation, organ damage and anemia, when sustained over time heme-driven repression of HGP can progress towards hypoglycemia, compromising host energy expenditure and thermoregulation. This hypometabolic state arrests the development of asexual stages of *Plasmodium spp*., which undergo pyknosis and develop mitochondrial dysfunction. In response, *Plasmodium* activates a transcriptional program reducing its virulence and inducing sexual differentiation towards the production of transmissible gametocytes. We infer that malaria-associated hypoglycemia represents a trade-off of an evolutionarily conserved defense strategy restricting *Plasmodium spp*. from accessing host-derived glucose and balancing parasite virulence and transmission.

## INTRODUCTION

Severe malaria refers to life-threatening clinical presentations of *P. falciparum* infection (Karnad et al., 2018; Marsh et al., 1995; Miller et al., 2002), in which hypoglycemia is a major independent risk factor of mortality (Madrid et al., 2015; Marsh et al., 1995; Service, 1995; White et al., 1983). The pathogenesis of malaria-associated hypoglycemia is thought to develop from a combination of events: anorexia of infection, an evolutionarily conserved component of sickness behavior that reduces appetite and nutrient intake from diet (Clark et al., 2008; Hart, 1988), as well as deregulation of hepatic glucose production (HGP)(Geoffrion et al., 1985; Kawo et al., 1990; Thien et al., 2006) and glucose consumption by *Plasmodium spp*. (Olszewski and Llinas, 2011; Preuss et al., 2012b; Thien et al., 2006) and immune cells (Buck et al., 2017; Man et al., 2017; Wang et al., 2019).

Whether malaria-associated hypoglycemia is a pathologic event *per se* or whether its pathologic effects represent a trade-off (Stearns and Medzhitov, 2015) of an otherwise adaptive response to *Plasmodium spp*. infection is unclear. Supporting its intrinsic pathologic nature, glucose administration to *Plasmodium chabaudi chabaudi (Pcc*)-infected mice is protective, while pharmacologic inhibition of glycolysis is detrimental (Cumnock et al., 2018). However, the reduction of blood glucose levels associated with caloric restriction (Mancio-Silva et al., 2017) or inhibition of glycolysis (Wang et al., 2018), are protective against *P. berghei* ANKA-infection in mice. Moreover, *P. falciparum* proliferation *in vitro* depends on glucose (Humeida et al., 2011) and reducing glucose concentration induces parasite pyknosis (Babbitt et al., 2012).

Malaria-associated hypoglycemia develops during the circadian cycles of parasite red blood cell (RBC) invasion, proliferation and hemolysis. Labile heme, an alarmin generated as a byproduct of hemolysis (Soares and Bozza, 2016) during the blood stage of *Plasmodium spp*. infection (Ademolue et al., 2017; Cruz et al., 2018; Elphinstone et al., 2016; Gouveia et al., 2017; Ramos et al., 2019) promotes the pathogenesis of severe malaria (Ferreira et al., 2008; Knackstedt et al., 2019; Pamplona et al., 2007). The pathologic effects of labile heme are countered by a host stress-responsive transcriptional program whereby the nuclear factor erythroid 2-related factor 2 (NRF2) induces the heme catabolizing enzyme, heme oxygenase-1 (HO-1) (Ferreira et al., 2011; Pamplona et al., 2007; Ramos et al., 2019; Seixas et al., 2009). While protective against malaria, heme catabolism by HO-1 does not appear to impose a fitness cost on the development of asexual stages of *Plasmodium spp*., establishing disease tolerance to malaria (Ferreira et al., 2011; Ramos et al., 2019). This defense strategy underlies the protective effect of sickle hemoglobin (Hb) against severe malaria (Ferreira et al., 2011).

Heme catabolism by HO-1 produces catalytic iron that can repress the transcription of the hepatic gluconeogenic enzyme glucose-6-phosphatase catalytic subunit 1 (*G6pc1*) (Weis et al., 2017). Given the excessive amounts of labile heme that accumulate in plasma during *Plasmodium* spp. infection in mice (Gouveia et al., 2017; Ramos et al., 2019) and humans (Ademolue et al., 2017; Cruz et al., 2018; Elphinstone et al., 2016; Knackstedt et al., 2019), we hypothesized that labile heme might contribute to the development of malaria-associated hypoglycemia. Here, we demonstrate that labile heme is sufficient *per se* to recapitulate the development of anorexia, repression of HGP and reduction of glycemia, observed in response to *Pcc* infection in mice. While protective against immune-mediated liver damage and anemia, sustained repression of HGP led to hypoglycemia, compromising the energetic demands of the infected host. This hypometabolic state is sensed by asexual stages of *Plasmodium*, which activate a transcriptional program associated with arrested development, reduction of virulence and sexual differentiation towards the production of transmissible gametocytes. These findings suggest that while labile heme elicits a protective hypometabolic response that imposes a fitness cost to *Plasmodium*, this defense strategy carries as a major trade-off the development of malaria-associated hypoglycemia, a defining feature of severe malaria.

## RESULTS

### HGP controls glycemia during *P**lasmodium* infection (*Fig. 1&S1*)

Intraperitoneal (i.p.) inoculation of *Pcc* AS-infected RBC (iRBCs) to C57BL/6 mice was characterized by increasing numbers of circulating iRBC during the first 7 days (*i*.*e*. peak of infection), decreasing thereafter (*Fig. 1A*). This was associated with the development of severe anemia (*Fig. 1A*), anorexia and fasting during the peak of infection (*Fig. 1B*)(Cumnock et al., 2018; Elased and Playfair, 1994). Moreover, there was a marked reduction of blood glucose concentration (*Fig. 1C*), paralleled by the repression of hepatic *G6pc1* mRNA (*Fig. 1D*) and protein (*Fig. 1E*) expression.

**Figure 1.**
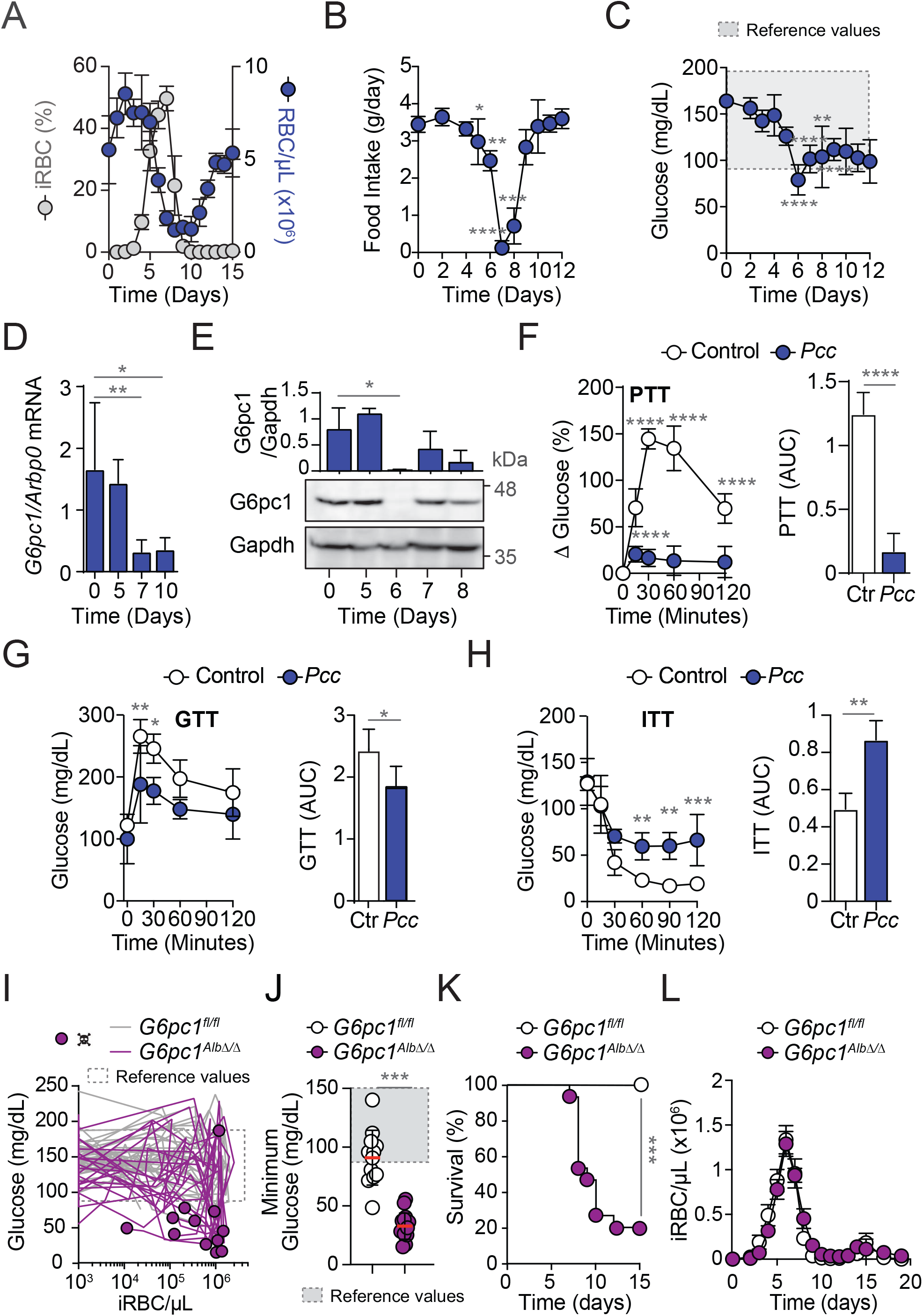
HGP controls glycemia during *Pcc* infection (*see also Figure S1*) (A) Parasitemia (% iRBC) and number of RBC *per* µL of blood in *Pcc-*infected C57BL/6 mice (N=6). Data represented as mean ± SD, pooled from two independent experiments with similar trend. (B) Daily food intake (mean ± SD) in *Pcc-*infected C57BL/6 mice (N=6). Data pooled from two independent experiments with similar trend. (C) Glycemia (mean ± SD) in same mice as (A). (D) Hepatic *G6pc1* mRNA (normalized to *Arbp0;* mean ± SD) from *Pcc*-infected C57BL6 mice (N=5-10 *per* day), quantified by qRT-PCR. Data pooled from 2 independent experiments with similar trend. (E) G6pc1 protein (normalized to Gapdh; mean ± SD) in same mice as (D), quantified by western blot (lower panel). (F) Percent variation (Δ) of glycemia (right panel) after pyruvate administration (PTT) to C57BL6 mice, naïve (Control; Ctr.; N=4) or at the peak of *Pcc* infection (N=4). Area under the curve (AUC; right panel) corresponding to PTT. Data from one experiment shown as mean ± SD. (G) Glycemia (right panel) after glucose administration (GTT) to C57BL6 mice, naïve (Control; Ctr.; N=5) or at the peak of *Pcc* infection (N=4). Area under the curve (AUC; right panel) corresponding to GTT. Data from one experiment shown as mean ± SD. (H) Glucose concentration (right panel) after insulin administration (ITT) to C57BL6 mice, naïve (Control; Ctr.; N=5) or at the peak of *Pcc* infection (N=4). Area under the curve (AUC; right panel) corresponding to ITT. Data from one experiment shown as mean ± SD. (I) Individual disease trajectories of *G6pc1*^*Alb*Δ/Δ^ (N=15; purple lines) and *G6pC1*^*fl/lfl*^ (N=12; gray lines) mice, established by the relationship of blood glucose concentration and parasite density throughout a 15 day course of *Pcc* infection. Data pooled from 3 independent experiments with similar trend. Circles indicate death of individual mice. (J) Minimum blood glucose concentration (mean ± SEM), throughout the 15-day course of *Pcc* infection in the same mice as (I). (K) Survival after *Pcc* infection, in the same mice as (I). (L) Pathogen load (i.e. number of iRBC *per* μL of blood; mean ± SEM) in the same mice as (I). *P* values calculated by Mann-Whitney U test in (B, C, J), One-Way ANOVA with Holm-Sidak’s multiple comparison test in (D, E), Two-Way ANOVA with Tukey’s multiple comparison test in (F-H, L), t test for AUC in (F-H) and Log-rank (Mantel-Cox) test (conservative) in (K). *ns*: non-significant; * *P*<0.05; ** *P*<0.01; *** *P*<0.001; **** *P*<0.0001.

Administration of a pyruvate bolus (*i*.*e*. pyruvate tolerance test; PTT) to C57BL/6 mice at the peak of *Pcc* infection failed to increase glycemia to the level observed in control, fasted mice (*Fig. 1F; S1A*). Pyruvate is a preferential hepatic gluconeogenic substrate (Hughey et al., 2014), suggesting that *Pcc* infection is associated with repression of HGP, as in human malaria (Thien et al., 2006).

Administration of a glucose bolus (*i*.*e*. glucose tolerance test; GTT) to C57BL/6 mice at the peak of *Pcc* infection failed to increase blood glucose levels in infected mice to the levels observed in control, fasted mice (*Fig. 1G*). This is in line with an overall increase in glucose consumption, as in human malaria (Madrid et al., 2017; Thien et al., 2006; White et al., 1983).

Administration of an insulin bolus (*i*.*e*. insulin tolerance test; ITT) to C57BL/6 mice at the peak of *Pcc* infection reduced glycemia to a lower extent, compared to control fasted mice (*Fig. 1H*). Insulin administration to control fasted mice caused seizures, not observed in *Pcc*-infected mice (*Fig. S1B*). This suggests that *Pcc-*infected mice develop insulin resistance as a protective mechanism against “grand mal” seizures (Arieff et al., 1974).

To determine whether HGP controls glycemia during *Pcc* infection (*Fig. 1D*), hepatic *G6pc1* was deleted in an inducible manner in *G6pc1*^*Alb*Δ/Δ^ mice (Mutel et al., 2011). Individual disease trajectories (Schneider, 2011), established by the relationship of glycemia and pathogen load, showed that *Pcc*-infected *G6pc1*^*Alb*Δ/Δ^ mice developed lethal hypoglycemia (*Fig. 1I-K; S1C*), under similar pathogen loads to control *G6pc1*^*fl/fl*^ mice (*Fig. 1L*). This suggests that HGP plays a central role in the regulation of glycemia during *Pcc* infection. Of note, insulin concentration in plasma was barely detectable at the peak of *Pcc*-infection in *G6pc1*^*Alb*Δ/Δ^ or *G6pc1*^*fl/fl*^ mice (*Fig. S1D*), suggesting that insulin does not play a central role in the regulation of glycemia during *Pcc* infection.

The individual disease trajectories established by the relationship of core body temperature *vs*. pathogen load, revealed that the decrease in core body temperature was more pronounced in *Pcc*-infected *G6pc1*^*Alb*Δ/Δ^ *vs. G6pc1*^*fl/fl*^ mice (*Fig. S1E-G*). This was not the case for body weight loss (*Fig. S1H-J*), suggesting that HGP supports thermoregulation, while promoting body weight loss during *Pcc* infection.

### HGP controls host Energy Expenditure during *P**cc* infection (*Fig. 2&S2*)

Sustained repression of hepatic *G6pc1* expression in *G6pc1*^*Alb*Δ/Δ^ mice was associated with a more pronounced reduction of energy expenditure (EE) in response to *Pcc* infection, compared to *G6pc1*^*fl/fl*^ mice (*Fig. 2A,B*). This could not be attributed to differences in daily food intake (*Fig. 2C*), which was reduced to the same extent in *Pcc*-infected *G6pc1*^*Alb*Δ/Δ^ *vs. G6pc1*^*fl/fl*^ mice (mg/day range), compared to non-infected genotype matched controls (*Fig. S2D*; *i*.*e*. g/day). Reduction of EE was instead associated with lower activity (*i*.*e*. lethargy) (*Fig. 2D; S2A*) and respiratory exchange ratio (RER), in *Pcc*-infected *G6pc1*^*Alb*Δ/Δ^ *vs. G6pc1*^*fl/fl*^ mice (*Fig. 2E,F*). The later was attributed to a decrease in the ratio of CO_2_ production (VCO_2_) (*Fig. 2G*) *vs*. O_2_ consumption (VO_2_) (*Fig. 2H*). This suggests that repression of HGP in response to *Pcc* infection delineates the lower viable threshold of host glycemia without compromising vital metabolic processes. Of note, steady-state EE (*Fig. S2B,C*), food intake (*Fig. S2D*), RER (*Fig. S2E,F*), VCO_2_ (*Fig. S2G*) and VO_2_ (*Fig. S2H*) were similar in *G6pc1*^*Alb*Δ/Δ^ *vs. G6pc1*^*fl/fl*^ mice.

**Figure 2.**
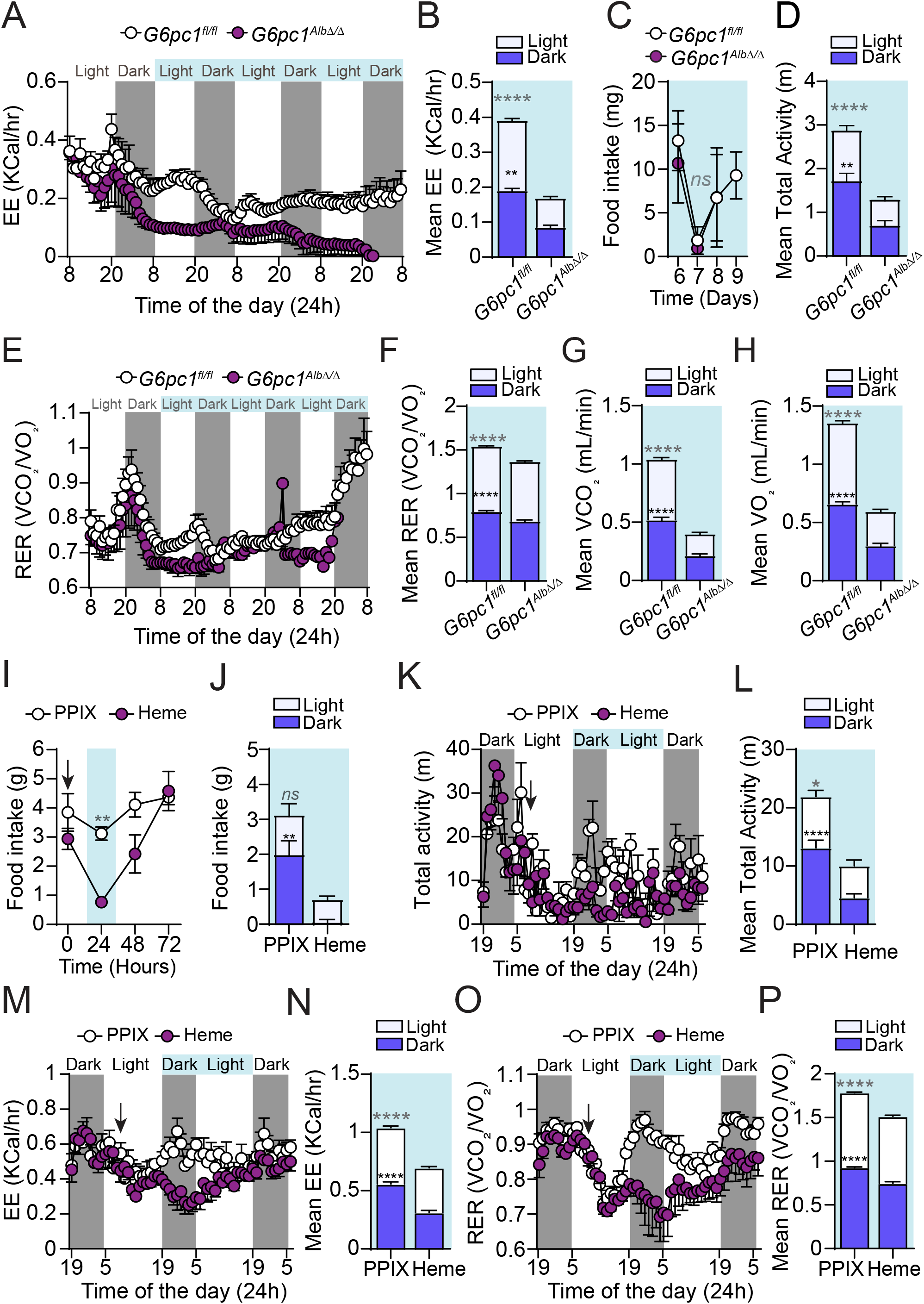
HGP regulates EE during *Plasmodium* infection (*see also Figure S2*) (A-H) Synchronized metabolic and behavioral quantification of *G6pc1*^*Alb*Δ/Δ^ (N=4; purple circles) and *G6pC1*^*fl/lfl*^ (N=4; white circles) mice, from 6 to 9 days after *Pcc* infection. Data from one experiment. (A) Hourly energy expenditure (EE), represented as mean ± SEM. (B) Mean of hourly EE (± SEM), corresponding to the light blue period in (A). Data segregated into daily light/dark cycle. (C) Mean daily food intake corresponding to the period indicated by light blue in (A) ± SEM, segregated into daily light/dark cycle. (D) Mean of total activity (1-hour binning values *per* mouse) corresponding to the period indicated by light blue in (A), ± SEM, segregated into daily light/dark cycle. (E) Hourly respiratory exchange ratio (RER) represented as mean ± SEM. (F) Mean of hourly RER corresponding to the period indicated by light blue in (E) ± SEM, segregated into daily light/dark cycle. (G) Mean of hourly volume of exhaled CO_2_ corresponding to the period indicated by light blue in (E) ± SEM, segregated into daily light/dark cycle. (H) Mean of hourly volume of inhaled O_2_ corresponding to the period indicated by light blue in (E) ± SEM, segregated into daily light/dark cycle. (I-P) Synchronized quantification of metabolic and behavioral parameters in C57BL/6 mice receiving heme (30 mg/Kg; N=4; brown circles) or control protoporphyrin IX (PPIX; N=4; white circles), at the times indicated by an arrow. Data from two experiment representative of three independent experiments with similar trend. (I) Mean daily food-intake ± SEM. (J) Mean food-intake corresponding to the 24h period highlighted in light blue in (I) ± SEM, segregated into daily light/dark cycle. (K) Total activity represented as mean ± SEM. (L) Mean total activity, corresponding to the 24h period highlighted in light blue in (I) ± SEM, segregated into daily light/dark cycle. (M) Energy expenditure (EE) represented as mean ± SEM. (N) Mean EE, corresponding to the 24h period highlighted in light blue in (I) ± SEM, segregated into daily light/dark cycle. (O) Hourly respiratory exchange ratio (RER), represented as mean ± SEM. (P) Mean EE, corresponding to the 24h period highlighted in light blue in (I) ± SEM, segregated into daily light/dark cycle. *P* values in all panels were determined using Two-Way ANOVA with Sidak’s multiple comparison test. *ns* – non-significant; ** - p<0.01; *** - p<0.001; **** - p<0.0001.

### Labile heme recapitulates the hypometabolic response elicited by *P**cc* infection (*Fig. 2&S2*)

We asked whether the accumulation of labile heme contributes to the hypometabolic response to malaria. Heme administration to naïve C57BL/6 mice recapitulated the hallmarks of sickness behavior associated with *Pcc* infection, including anorexia (*Fig. 2I,J*) and lethargy (*Fig. 2K,L; S2I,J*). Other components of the hypometabolic response to *Pcc* infection, including reduced EE (*Fig. 2M,N*) and RER (*Fig. 2O,P*), with the latter reflecting a decrease of CO_2_ production (VCO_2_) *vs*. O_2_ consumption (VO_2_) (*Fig. S2K-N*), were also induced in response to heme administration. Protoporphyrin IX (PPIX) lacking the iron atom present in heme (*i*.*e*. Fe-protoporphyrin IX), did not induce any of these changes (*Fig. 2I-P; S2I-N*).

Consistent with labile heme repressing hepatic *G6Pc1 in vivo* (Weis et al., 2017), heme administration to fasted mice reduced the transient increase in glycemia in a PTT (*Fig. S2O*). This is in agreement with labile heme repressing HGP *in vivo*, similar to *Pcc* infection.

### Labile heme recapitulates the repression of hepatic gene expression in response to *P**cc* infection (*Fig. 3 & S3*)

Considering the liver as a metabolic hub, beyond HGP, we asked whether labile heme exerts additional effects, contributing to the state of negative energy balance imposed by malaria. For that purpose, we analyzed bulk RNA-sequencing data from the livers of mice infected with *Pcc* or receiving heme. Fasted mice were used as controls to account for the development of anorexia in *Pcc*-infected and heme-treated mice. Principal component analysis (PCA) showed a marked difference in the hepatic transcriptomic profiles of *Pcc*-infected *vs*. heme-treated mice, compared to control fasted mice, respectively (*Fig. S3A*). Unsupervised hierarchical clustering segregated the differentially expressed transcripts (*i*.*e*. 9,623) into four clusters. These included transcripts induced preferentially upon *Pcc* infection (Cluster 1) or heme administration (Cluster 2), as well as transcripts repressed preferentially by *Pcc* infection (Cluster 3) or heme administration (Cluster 4), as compared to control fasted mice, respectively (*Fig. 3A*).

**Figure 3.**
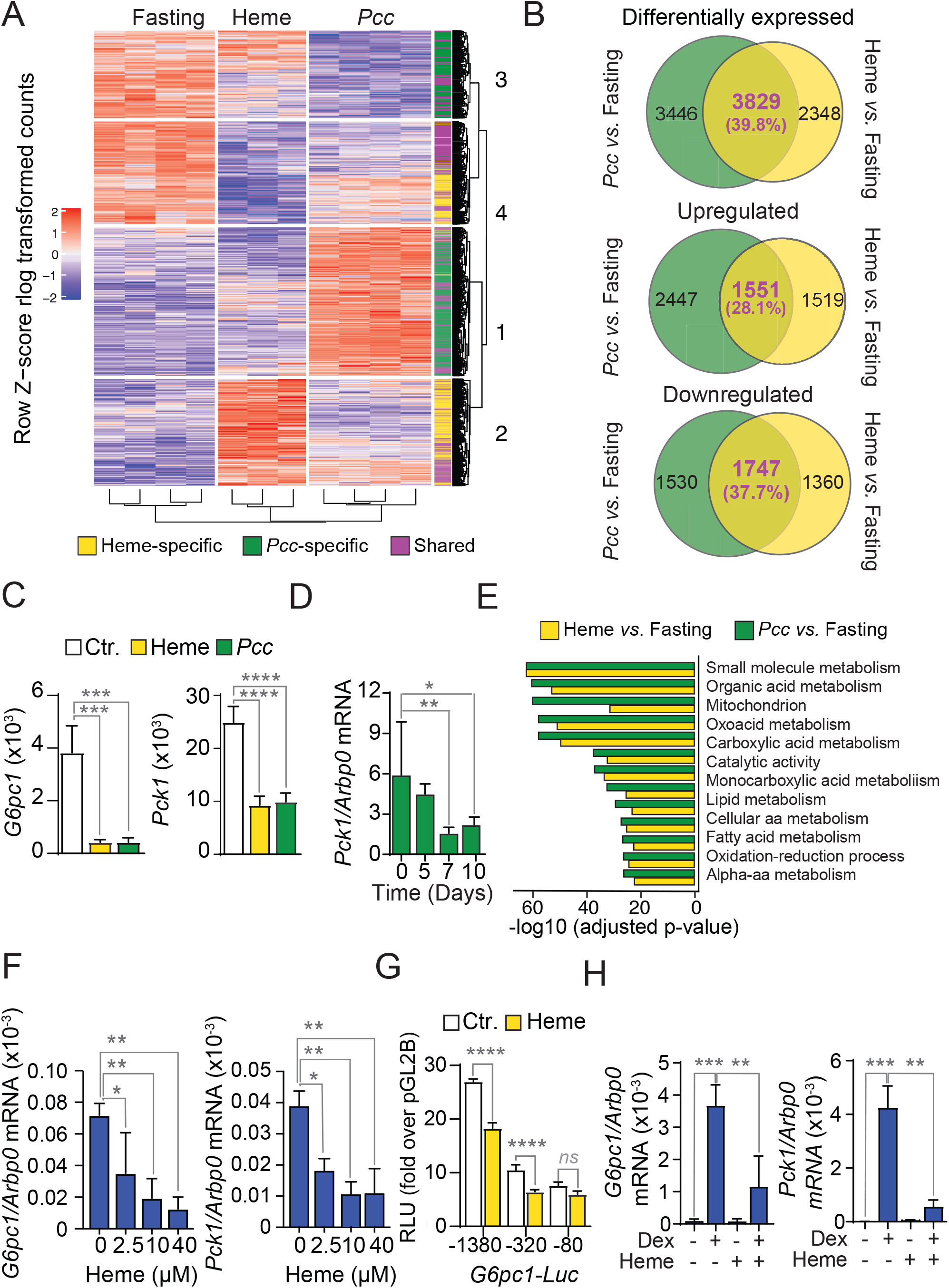
Labile heme recapitulates the repression of HGP in response to *Plasmodium* infection. (*see also Figure S3*) (A-C) Bulk RNA-seq analyzes from the liver of C57BL/6 mice subjected to fasting (Overnight; 15h; N=4), receiving heme (30mg/Kg, Overnight; N=3) or infected with *Pcc* (at the peak of infection; N=4) (A) Heatmap of unsupervised clustering analysis of 18,163 differentially expressed Z-score log transformed transcripts. Purple: Transcripts with common pattern of expression in mice receiving heme and infected with *Pcc*, compared to fasted control mice, respectively. Green: Transcripts with unique expression pattern in mice receiving heme. Yellow: Transcripts with an expression pattern unique to *Pcc* infected mice. (B) Venn diagrams of transcripts differentially expressed (*top panel*), transcripts upregulated (*middle panel*) or transcripts down regulated (*bottom panel*), in mice receiving heme or *Pcc*-infected mice, compared with control fasted mice, respectively. (C) Normalized *G6pc1* and *Pck1* read counts from RNA-seq in (A). (D) Hepatic *Pck1* mRNA expression (mean ± SD), in *Pcc*-infected C57BL6 mice (N=5-10 *per* day), quantified by qRT-PCR and normalized to *Arbp0*. Data pooled from 2 independent experiments with similar trend. (E) Functional enrichment analysis, using GO terms database, of transcripts down regulated in mice receiving heme and *Pcc*-infected mice. Only pathways with the highest enrichment score (−log10 adjusted p-value) are represented. (aa: amino acid). (F) *G6pc1* (*left panel*) and *Pck1* (*right panel*) mRNA expression (mean ± SD) in primary mouse hepatocytes exposed *in vitro* to heme (12h), quantified by qRT-PCR and normalized to *Arbp0*. Data from one out of 3 independent experiments with similar trend. (G) Relative luciferase units (RLU) in HepG2 cells transiently co-transfected *in vitro* with firefly luciferase reporters, regulated by the full length (−1380) or truncated (−320 and -80) *G6pc1* promoters and a Renilla luciferase reporter regulated by a minimal CMV promoter. Firefly RLU were normalized to Renilla Luciferase activity and data is represented as fold induction, compared to HepG2 cells transiently transfected with an empty firefly luciferase reporter (pGL2B), (*i*.*e*. baseline; 1 RLU). HepG2 cells were exposed to heme (40µM) 48h after transfection and analyzed 12h thereafter. Data represented as mean ± SD, pooled from 2 independent experiments with similar trend. (H) *G6pc1* (*left panel*) *Pck1* (*right panel*) mRNA expression (mean ± SD) in primary mouse hepatocytes exposed to Dexamethasone (Dex; 100 nM), heme (10µM) or Dexamethasone plus heme (12h), quantified by qRT-PCR and normalized to *Arbp0*. Data pooled from 2 independent experiments with similar trend. *P* values in (C,D,F-H) determined using One-Way ANOVA with Sidak’s multiple comparison test. Hypergeometric p-values in (E) were computed after correction for multiple testing. *ns* – non-significant; *p<0.05; **p<0.01; ***p<0.001; ****p<0.0001.

There was approximately 40% overlap in the identity of the differentially expressed hepatic transcripts in mice infected by *Pcc* or receiving heme, as compared to control fasted mice, respectively (*Fig. 3B; S3B*). This was attributed to a 28.1% and 37.7% overlap in induced or repressed transcripts, respectively (*Fig. 3B*). *Hmox1*, encoding HO-1, was among the overlapping induced transcripts (*Fig. S3B*), while *G6pc1* and *Pck1*, encoding phosphoenolpyruvate carboxykinase-cytosolic form (Pepck-c) (*Fig. 3C; S3B*) were among the overlapping repressed transcripts. Repression of the latter during *Pcc* infection was confirmed by qRT-PCR (*Fig. 3D*; See also *Fig. 1D*). This suggests that labile heme orchestrates a hepatic transcriptional response underlying the hypometabolic state imposed by *Plasmodium* infection.

### Labile heme mimics the repression of HGP during malaria (*Fig. 3 &S3*)

Functional enrichment analysis showed an overlap of the 12 metabolic pathways most significantly repressed in response to *Pcc* or heme administration, as compared to control fasted mice, respectively (*Fig. 3E; Table S1*). These were involved in “regulation of small molecule metabolism”, which includes gluconeogenesis, organic acid, oxoacid, carboxylic acid, monocarboxylic acid, lipid and amino acid metabolism (*Fig. 3E*). A broader analysis showed a 53% overlap of all pathways repressed in response to *Pcc* infection or heme administration (*Fig. S3C*).

In contrast, the pathways most significantly induced in response to *Pcc* infection (*i*.*e*. immune and stress responses to biotic and/or organic substances) diverged from those induced upon heme administration (*i*.*e*. subcellular processes)(*Fig. S3D,E; Table S2*). When accounting for all pathways induced by *Pcc* infection, only 23% were also induced in response to heme administration (*Fig. S3F; Table S2*).

The regulatory regions of the overlapping genes repressed during *Pcc* infection and in response to heme administration, shared a number of GC-rich DNA-binding motifs recognized by the SP/KLF (Specificity protein/Kruppel-like factor) family of transcriptional regulators, including SP1, SP2, KLF3, KLF5, KLF7 and ZF5 (*Fig. S3G*). The regulatory regions of these genes also shared DNA-binding motifs recognized by the E2F-1, E2F-3 and E2F-4 members of the E2F family of transcriptional regulators as well as by the zinc finger transcriptional regulators early growth response protein 1 (EGR1/KROX), CTCF or WT1 and by the Aryl hydrocarbon receptor (AhR)(*Fig. S3G*). A broader analysis showed an additional number of shared but also non-shared (*Fig. S3G*) DNA-binding motifs in the regulatory regions of ∼60% of these genes (*Fig. S3H; Table S3*).

In keeping with labile heme repressing *G6pc1* transcription (*Fig. 1D*)(Yin et al., 2007), *G6pc1* mRNA expression by primary mouse hepatocytes was repressed, in a dose-dependent manner, by exogenous heme, *i*.*e*. labile heme (*Fig. 3F*). To explore further how heme represses *G6pc1* transcription, we used human HepG2 cells transiently transfected *in vitro* with luciferase reporters (*Fig. 3G*). Heme inhibited to the same extent (*i*.*e*. 31-36%) the transcription of *G6pc1*-luciferase reporters regulated by the -1380/+60 or the -320/+60 base pair (bp) promoter region (*i*.*e*. from the TATA box), as compared to vehicle-treated control cells, respectively (*Fig. 3G*). In contrast, heme failed to repress the transcription of a minimal *G6pc1*-luciferase reporter controlled by the -80/+60 bp promoter region (*Fig. 3G*). This suggests that labile heme represses *G6pc1* transcription via a mechanism that targets one or several transcriptional regulators acting at the -320 to -80 bp region of *G6pc1* promoter. This promoter region contains DNA-binding motifs recognized by SP1, SP2, KLF3, KLF5, EGR1 and Ahr but not by CTCF or WT1 (*Fig. S3I*), suggesting that labile heme represses hepatic *G6pc1* transcription via a mechanism targeting one or several members of the SP/KLF family of transcription regulators and/or Ahr.

Glucocorticoids induce hepatic *G6pc1* transcription via SP/KLF family of transcriptional regulators (Cui et al., 2019) and prevent the development of malaria-associated hypoglycemia (Vandermosten et al., 2018). Labile heme repressed the induction of *G6pc1* and *Pck1* mRNA in primary mouse hepatocytes exposed to the synthetic glucocorticoid dexamethasone (*Fig. 3H*), suggesting that labile heme impairs the protective effects of glucocorticoids against malaria-associated hypoglycemia.

### Repression of HGP restrains immune activation and is protective against liver damage (*Fig. 4, S4*)

Immune activation relies to a large extent on glycolysis, required to fuel anabolic metabolism and leukocyte proliferation (Pearce et al., 2013). In support of the idea that regulation of HGP controls the immune response to *Plasmodium* infection, there was a marked reduction in the number of splenic natural killer (NK) and natural killer T (NKT) cells in *Pcc*-infected *G6pc1*^*Alb*Δ/Δ^ *vs. G6pc1*^*fl/fl*^ mice (*Fig. 4A,B*). Splenic dendritic cell (*Fig. S4A,B*), monocyte (*Fig. S4A,C*) and red pulp macrophage (*Fig. S4A,D*) numbers were similar, while the number of circulating neutrophils was higher in *Pcc*-infected *G6pc1*^*Alb*Δ/Δ^ *vs. G6pc1*^*fl/fl*^ mice (*Fig. S4E*).

**Figure 4.**
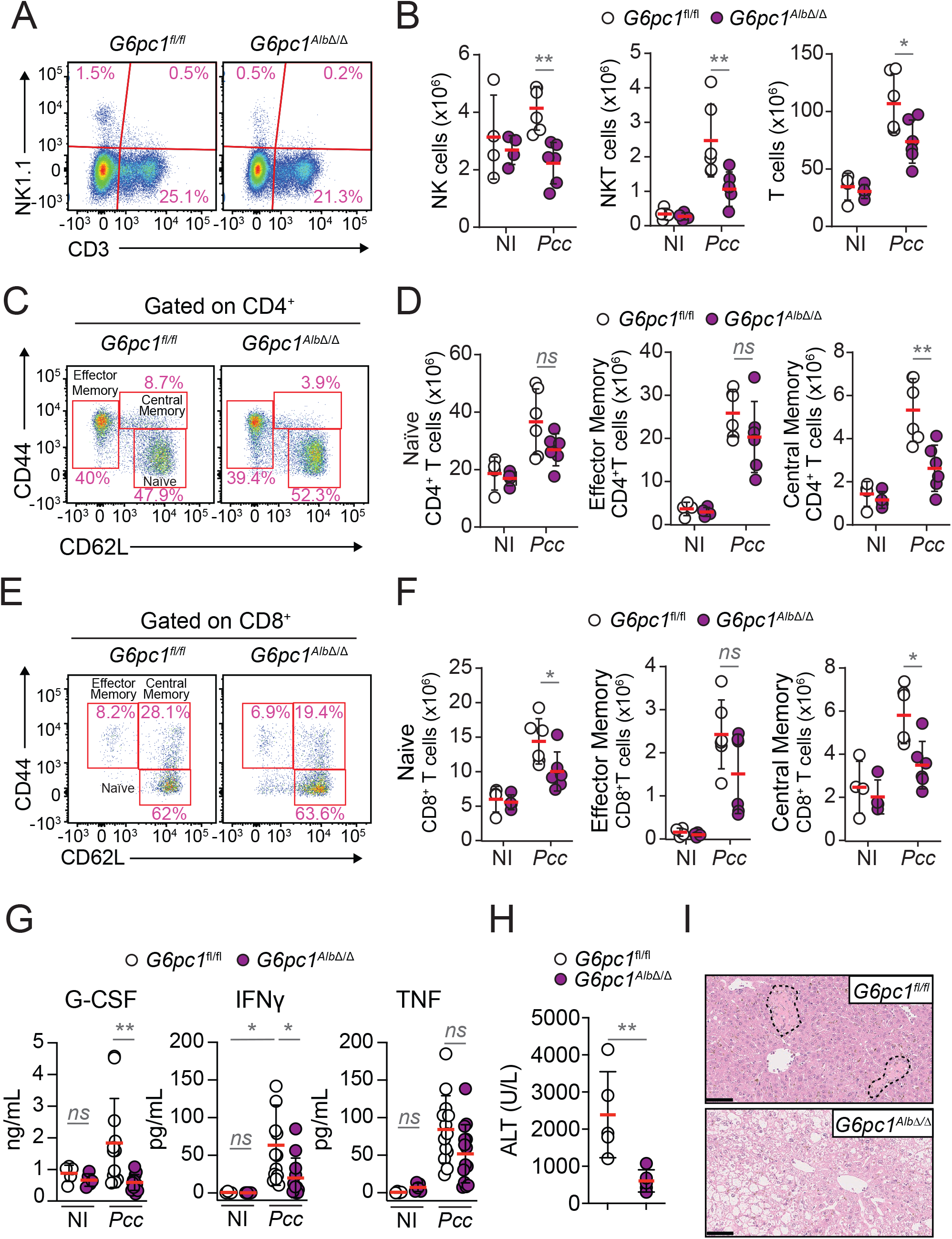
Repression of HGP restrains immune-driven inflammation and liver damage in response to *Pcc* infection. (*see also Figure S4*) (A, B) Splenic NK (NK1.1^+^CD3^-^), NKT (NK1.1^+^CD3^+^) and T (NK1.1^-^CD3^+^) cells in *G6pc1*^*Alb*Δ/Δ^ (purple dots) and *G6pC1*^*fl/lfl*^ (white dots) mice not infected (NI; N=4 *per* genotype) and at the peak of *Pcc*-infection (N=6 per genotype). Data pooled from 2 independent experiments with similar trend. (A) Representative FACS plots. (B) Quantification of (A) represented as mean (red bar) ± SD, from individual mice represented by circles. (C, D) Splenic naïve (CD4^+^CD44^low^CD62L^+^), effector memory (CD4^+^CD44^+^CD62L^-^) and central memory (CD4^+^CD44^+^CD62L^+^) T helper cells in the same mice as (A). (C) Representative FACS plots for CD44 and CD62L staining, gated on CD4^+^ cells. (D) Quantification of (C) represented as mean (red bar) ± SD, from individual mice represented by circles. (E, F) Splenic naïve (CD8^+^CD44^low^CD62L^+^), effector memory (CD8^+^CD44^+^CD62L^-^) and central memory (CD8^+^CD44^+^CD62L^+^) CD8^+^ T cells in the same mice as (A). (E) Representative FACS plots. (F) Quantification of (E) represented as mean (red bar) ± SD, from individual mice represented by circles. (G) G-CSF, IFNγ and TNF concentrations in plasma of non-infected (NI) and at the peak (day 7) of *Pcc* infection in *G6pc1*^*Alb*Δ/Δ^ (N=12; purple circles) and *G6pC1*^*fl/lfl*^ (N=11; white circles) mice. Data represented as mean (red bar) ± SD, from individual mice, pooled from 2 independent experiments with similar trend. (H) Alanine Aminotransferase (ALT) concentration in plasma of non-infected (NI) and at the peak (day 7) of *Pcc* infection in *G6pc1*^*Alb*Δ/Δ^ (N=5; purple circles) and *G6pC1*^*fl/lfl*^ (N=5; white circles) mice. Circles represent individual mice, red bars mean values ± SD, from one experiment. (I) H&E staining of the liver from *Pcc*-infected *G6pc1*^*Alb*Δ/Δ^ (N=7) and *G6pC1*^*fl/lfl*^ (N=5) mice. Dashed lines outline necrosis areas. Images are representative of 2 independent experiments. Scale bar: 100 μm. P values in (B,D,F,G) determined using Two-Way ANOVA with Tukey’s multiple comparison test and in (H) using the Mann Whitney test. *ns* – non-significant; * - p<0.05; ** - p<0.01.

The number of splenic T cells was also reduced in *G6pc1*^*Alb*Δ/Δ^ *vs. G6pc1*^*fl/fl*^ mice (*Fig. 4A,B*). This was associated with a reduction of splenic central memory (CD4^+^CD44^+^CD62L^+^) T helper (T_H_) cells (*Fig. 4C,D*) but not naïve (CD4^+^CD44^Low^CD62L^+^) or effector memory (CD4^+^CD44^+^CD62L^-^) T_H_ cells (*Fig. 4C,D*). The number of splenic central memory (CD8^+^CD44^+^CD62L^+^) cytotoxic T cells and naïve (CD8^+^CD44^low^CD62L^+^) cytotoxic T cells was also reduced in *Pcc* infected *G6pc1*^*Alb*Δ/Δ^ *vs. G6pc1*^*fl/fl*^ mice (*Fig. 4E,F*). This was not the case for effector memory (CD8^+^CD44^+^CD62L^-^) cytotoxic T cells (*Fig. 4E,F*). Taken together, these data suggest that repression of HGP restrains the activation and/or expansion of central memory T_H_ and cytotoxic T cells in response to *Plasmodium* infection.

*Pcc*-infected *G6pc1*^*Alb*Δ/Δ^ mice also showed a marked reduction in granulocyte-colony stimulating factor (G-CSF) and interferon gamma (IFNγ) concentration in plasma, compared to *G6pc1*^*fl/fl*^ mice (*Fig. 4G*). Other cytokines and chemokines were not affected, except for eotaxin (*Fig. S4F*) and TNF, but the later did not reach statistical significance (*Fig. 4G*).

Repression of HGP in *Pcc*-infected *G6pc1*^*Alb*Δ/Δ^ mice was associated with protection from liver damage, compared to *G6pc1*^*fl/fl*^ mice. This was revealed by a lower accumulation of alanine transaminase (ALT) in plasma (*Fig. 4H*) and confirmed by histological analyzes (*Fig. 4I*).

### Repression of HGP regulates *P**lasmodium* behavior (*Fig. 5&S5*)

*P. falciparum* development and proliferation *in vitro* are dependent on glucose (Hirako et al., 2018; MacRae et al., 2013; Olszewski and Llinas, 2011; Salcedo-Sora et al., 2014; Srivastava et al., 2016), suggesting that repression of HGP might impact on *Plasmodium spp*. development and/or proliferation *in vivo*. In support of this hypothesis, the number of circulating RBC was higher throughout the course of *Pcc* infection in *G6pc1*^*Alb*Δ/Δ^ *vs. G6pc1*^*fl/fl*^ mice (*Fig. 5A,B; S5A*). In contrast, the maximal percentage of circulating iRBC was lower in *G6pc1*^*Alb*Δ/Δ^ *vs. G6pc1*^*fl/fl*^ mice, “normalizing” the pathogen burden among genotypes (*Fig. 5A,B*).

**Figure 5.**
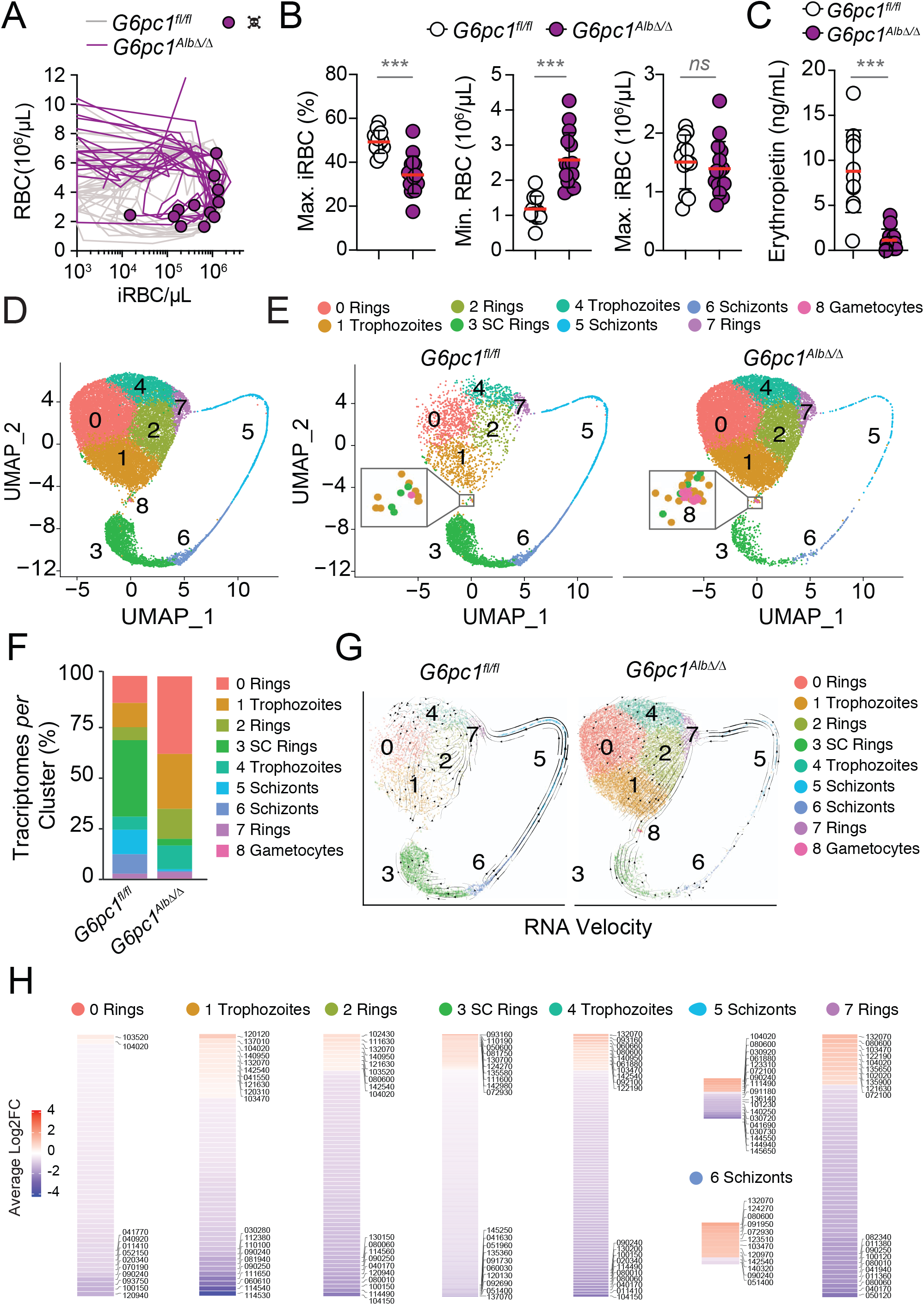
Repression of HGP regulates *Plasmodium* behavior (*see also Figure S5*) (A) Disease trajectories established by the relationship between circulating RBC and infected RBC (iRBC) numbers per µL of blood, during the 15-day course of *Pcc* infection in *G6pc1*^*Alb*Δ/Δ^ (N=15; purple lines) and *G6pC1*^*fl/lfl*^ (N=12; gray lines) mice. Purple circles indicate death. (B) Maximum parasitemia (% iRBC), minimum RBC (*i*.*e*. anemia) and maximum iRBC (pathogen load) numbers *per* µL of blood, during the course of *Pcc* infection in the same mice as (A). Data is represented as mean (red bars) ± SEM, pooled from 3 independent experiments with similar trend. Circles represent individual mice. (C) Erythropoietin concentration (mean ± SD) in plasma from *G6pc1*^*Alb*Δ/Δ^ (N=12; purple circles) and *G6pC1*^*fl/lfl*^ (N=11; white circles) mice, at the peak of *Pcc* infection (*i*.*e*. day 7). Data from 2 independent experiments. Circles represent individual mice. (D) Combined Single-cell (sc) RNA UMAP projection of single-cell parasite transcriptomes in FACS-sorted iRBCs from *G6pc1*^*fl/lfl*^ and *G6pc1*^*Alb*Δ/Δ^ mice. Each dot represents an individual parasite transcriptome. Colors and corresponding cluster numbers identify parasite developmental stages, according to 25 highest expressed transcripts and analogy to *P. falciparum* and *P. berghei* gene orthologues (Malaria Cell Atlas; https://www.sanger.ac.uk/tool/mca/). (E) scRNA UMAP projection of parasite transcriptomes in the FACS-sorted iRBCs from *G6pc1*^*fl/lfl*^ (n= 5294; left panel) and *G6pc1*^*Alb*Δ/Δ^ (n= 15543 right panel) mice. (F) Percentage of *Pcc* transcriptomes per cluster in FACS-sorted iRBCs from (D). (G) RNA velocity analysis of *Pcc* transcriptomes in FACS-sorted iRBCs from (D). (H) Cluster-specific heatmap of differentially regulated genes (average log_2_fold change) in the same FACS-sorted iRBCs as (D). Numbers correspond to *Pcc* gene identification (e.g. PCHAS_xxxx). P values in (B, C) were determined using the Mann Whitney test. *ns*: non-significant; ***: p<0.001.

That repression of HGP limits the severity of anemia imposed by *Plasmodium* infection was corroborated by an increase of Hb concentration, hematocrit and mean corpuscular hemoglobin concentration (MCHC), as well as a decrease in RBC mean corpuscular volume (MCV) in *Pcc*-infected *G6pc1*^*Alb*Δ/Δ^ *vs. G6pc1*^*fl/fl*^ mice (*Fig. S5A*). RBC mean corpuscular hemoglobin (MCH) and distribution RBC width (RDW) were also closer to the physiological range in *Pcc*-infected *G6pc1*^*Alb*Δ/Δ^ compared to *G6pc1*^*fl/fl*^ mice (*Fig. S5A*).

Additionally, *Pcc*-infected *G6pc1*^*Alb*Δ/Δ^ also had a marked decrease in erythropoietin concentration in plasma, compared to *G6pc1*^*fl/fl*^ mice (*Fig. 5C*). Given that erythropoietin is the main hormone responsible for the induction of erythropoiesis in response to anemia, this observation supports further the idea that repression of HGP is protective against the anemia caused by *Plasmodium* infection.

We hypothesized that repression of HGP restrains anemia via a mechanism that acts on *Plasmodium*. To test this hypothesis we performed single-cell RNA-sequencing (scRNAseq) in (GFP^+^) iRBC FACS-sorted from *G6pc1*^*Alb*Δ/Δ^ and *G6pc1*^*fl/fl*^ mice. Parasite transcriptomes were segregated into nine clusters (*Fig. 5D,E*), ascribed to different developmental stages, according to the 25 most highly expressed transcripts and by comparison to *P. falciparum* and *P. berghei* gene orthologues (*Table S4*). Clusters 0, 2 and 7 (*Table S4*) corresponded to early ring stages (*Fig. 5E; S5B*) and clusters 1 and 4 to trophozoites (*Fig. 5E; S5B*). Cluster 3 included transcripts associated with ring stages, trophozoites and early schizont stages (*Table S4*), but also with *Plasmodium* sexual commitment, suggesting that cluster 3 includes early stages of gametocyte development (*Fig. 5E; S5B*). Clusters 5 and 6 (*Table S4*) corresponded to schizonts, while cluster 6 was associated with early schizonts (*Fig. 5E; S5B*). Cluster 8 corresponded to gametocytes (*Table S4*; *Fig. 5E; S5B*).

The number of parasite transcriptomes *per* iRBC from *G6pc1*^*Alb*Δ/Δ^ mice was 3-fold higher, compared to *G6pc1*^*fl/fl*^ mice (*Fig. 5E,F*). This was not related to the number of iRBC (GFP^+^) processed (*Fig. S5C*). The frequency of transcriptomes corresponding to each *Plasmodium* cluster was also different in iRBC from *G6pc1*^*Alb*Δ/Δ^ *vs. G6pc1*^*fl/fl*^ mice (*Fig. 5E,F*). There was a relative increase in the number of transcriptomes corresponding to ring stages and trophozoites (*i*.*e*. clusters 0-2, 4, 7, 3) and a reduction in transcriptomes corresponding to schizonts (*i*.*e*. clusters 5, 6). Moreover, there was an additional parasite transcriptome cluster in iRBC from *G6pc1*^*Alb*Δ/Δ^ mice, corresponding to gametocytes (*i*.*e*. cluster 8), which as barely detected in iRBC from *G6pc1*^*fl/fl*^ mice (*Fig. 5E,F*). This suggests that parasites from *G6pc1*^*Alb*Δ/Δ^ mice undergo gametocytogenesis (Yuda et al., 2015). Of note, transcripts corresponding to the parasite glycolytic enzymes were detected preferentially in clusters 1, 4, 5, 6, and 8, corresponding to trophozoites, schizonts and gametocytes, respectively. In contrast, transcripts corresponding to TCA-related enzymes were detected mainly in cluster 8, corresponding to gametocytes, and to a lower extent in clusters 1 and 6 (*Table S5*).

RNA velocity analysis of temporal transcriptome transitions (Bergen et al., 2020) suggested that parasites in *G6pc1*^*fl/fl*^ mice followed the expected developmental progression through schizonts (*i*.*e*. cluster 5/6) towards ring stages and trophozoites (*i*.*e*. clusters 0,1,2,4,7), with minimal sexual commitment (*i*.*e*. cluster 3, 8). This was severely altered in parasites from *G6pc1*^*Alb*Δ/Δ^ mice, in that schizonts (*i*.*e*. cluster 6) appear to undergo sexual commitment (*i*.*e*. cluster 3), as revealed by the expression of transcriptomes corresponding to gametocytes (*i*.*e*. cluster 8) (*Fig. 5G*). Moreover, there was an apparent “backward transition” from trophozoites (*i*.*e*. clusters 0,1,2,4) and schizonts (*i*.*e*. cluster 5) towards rings (*i*.*e*. cluster 7) as well as an arrest of schizont transition from cluster 5 to cluster 6 (*Fig. 5G*). This suggests that repression of HGP is associated with a developmental arrest of asexual stages of *Plasmodium*, while favoring sexual commitment. Of note, there was a marked repression of transcripts involved in DNA replication and transcriptional regulation (*Fig. 5H, Table S6*), while transcripts related to translation, but also heat-shock proteins, were induced in parasites from *G6pc1*^*Alb*Δ/Δ^ *vs. G6pc1*^*fl/fl*^ mice (*Fig. 5H, Table S6*). Functional enrichment analysis confirmed the repression of DNA replication (*Fig. S5D*) and anabolic metabolism (*Fig. S5E,F*).

### Repression of HGP reduces *P**lasmodium* virulence (*Fig. 5&6*)

Parasites from *G6pc1*^*Alb*Δ/Δ^ mice reduced the expression of transcripts corresponding to virulence factors (*Fig. 5H, Table S6*), as confirmed for *hmgb2* (Briquet et al., 2015) and *famb* (Brugat et al., 2017), by qRT-PCR (*Fig. 6A*). Infection of *G6pc1*^*Alb*Δ/Δ^ mice with parasites isolated from *G6pc1*^*Alb*Δ/Δ^ mice, led to lower incidence of mortality, compared to infection with parasites from *G6pc1*^*fl/fl*^ mice (*Fig. 6B*). This was associated with reduced temperature-loss (*Fig. S6A*), without changes in pathogen load (*Fig. 6B*) or glycemia (*Fig. S6B*). This data shows that repression of HGP reduces *Pcc* virulence.

**Figure 6.**
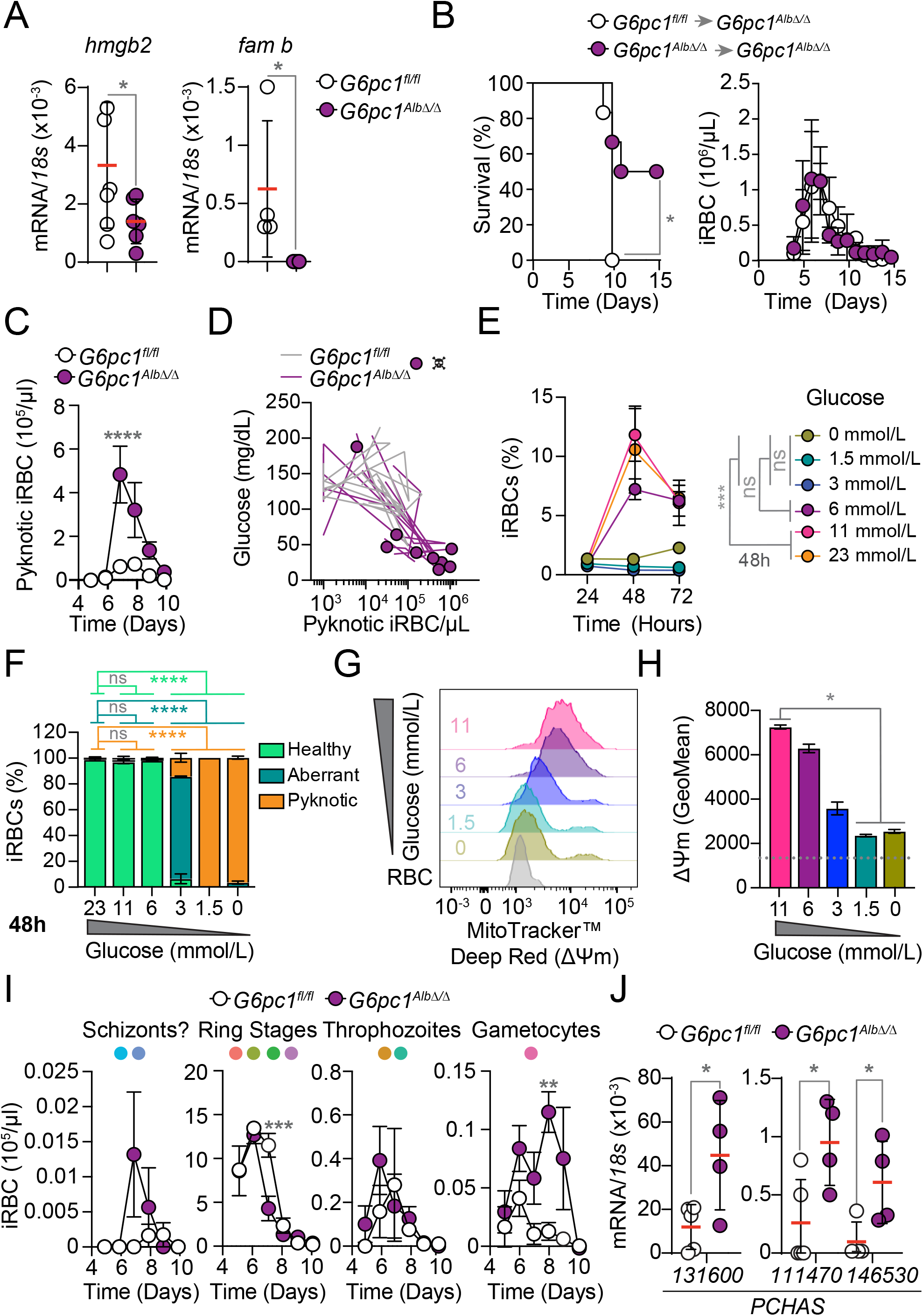
Repression of HGP regulates *Plasmodium* virulence and gametocytogenesis (*see also Figure S6*) (A) Quantification of hepatic mRNA encoding the virulence factors *hmgb2* (PCHAS_072200; N=6 *per* genotype) and *fam b* (PCHAS_100100; N=4 per genotype). Data is represented as mean (red bars) ± SD, pooled from 2 independent experiments with similar trend. Circles represent individual mice. Data for *hmgb2* and *famb* was analyzed 7 and 8 days after *Pcc* infection, respectively. (B) Survival (*left panel*) and pathogen load (*right panel;* mean ± SD) of *G6pc1*^*Alb*Δ/Δ^ mice infected with *Pcc* iRBC from *G6pc1*^*Alb*Δ/Δ^ (purple circles) *vs. G6pc1*^*fl/lfl*^ (light purple circles) mice (N=6 per group). Data pooled from 2 independent experiments with similar trend. (C) Number of iRBC (mean ± SD) containing pyknotic parasites, quantified from Giemsa-stained thin blood smears of *Pcc*-infected *G6pc1*^*Alb*Δ/Δ^ (N=10) and *G6pC1*^*fl/lfl*^ (N=8) mice. Data pooled from 2 independent experiments with similar trend. (D) Disease trajectories established by the relationship between number of iRBC containing pyknotic parasites, calculated from Giemsa-stained thin blood smears in the same mice as in (C) and glycemia, during the 15-day course of *Pcc*. Purple circles indicate death. (E-F) Glucose concentration regulates *P. falciparum* development and proliferation *in vitro*. (E) Percentage of *P. falciparum* infected human RBC (iRBC) cultured *in vitro* under varying glucose concentrations (*i*.*e*. from 23-0 mmol/L). Data calculated from Giemsa-stained thin blood smears and presented as mean ± SEM, from three wells in 1 out of 3 independent experiments with similar trend. (F) Percentage of *P. falciparum* iRBC containing morphologically healthy, aberrant or pyknotic parasites at 48h, from the same samples as in (E). Data is presented as mean ± SEM, from three wells in1 out of 3 independent experiments with similar trend. (G-H) Glucose regulates *P. falciparum* mitochondrial function *in vitro*. (G) Representative histograms of mitochondrial membrane potential (ΔΨm; MitoTracker™ Deep Red FM) in *P. falciparum*-infected human RBC (iRBC) cultured for 24h *in vitro* under varying glucose concentrations (*i*.*e*. from 11-0 mmol/L). *P. falciparum* iRBC were identified as GFP and Hoechst fluorescent cells (*See S6H*). Gey histogram represents background MitoTracker™ Deep Red FM staining in non-infected (Hoechst negative) RBC. Histograms are representative of 4 replicates, stained in duplicate, from one experiment. (H) Quantification of Geometric Mean Fluorescence Intensity (mean ± SEM) of MitoTracker™ Deep Red FM from (G). Dotted grey line indicates background signal in non-infected RBC, averaged from all replicates analyzed. (I) Quantification of different *Pcc* development stages (mean ± SD), identified morphologically from the same Giemsa-stained thin blood smears as (C). (J) Quantification of mRNA encoding gametocyte-specific genes in whole blood from *Pcc*-infected *G6pc1*^*Alb*Δ/Δ^ (N=4) vs. *G6pc1*^*fl/lfl*^ (N=4-5) mice, 9 days after *Pcc* infection, normalized to *Pcc 18S* mRNA. PCHAS_131600: Gamete egress and sporozoite traversal protein (*gest*), PCHAS_146530: Osmiophilic body protein *(g377*), PCHAS_111470: Gametogenesis essential protein 1 *(gep1*). Data is represented as mean (red bars) ± SD, pooled from 2 independent experiments with similar trend. Circles represent individual mice. P values were determined in (A,J) using One-tailed Unpaired Mann Whitney U test, in (B) using Log-rank (Mantel-Cox) test (conservative) for survival and Two Way ANOVA for pathogen load, in (C,I) using Two Way ANOVA with Sidak’s multiple comparison test for time course, in (E,F) using Two Way ANOVA with Tukey’s multiple comparison test and in (H) using Kruskal-Wallis tests with Dunn’s multiple comparison test. *ns* – non-significant; * - p<0.05; ** - p<0.01; *** - p<0.001; **** - p<0.0001.

### Repression of HGP induces *P**lasmodium* pyknosis (*Fig. 6&S6*)

The number of circulating iRBC containing pyknotic parasites was increased by 7.6-fold at the peak of *Pcc* infection in *G6pc1*^*Alb*Δ/Δ^ *vs. G6pc1*^*fl/fl*^ mice, as assessed morphologically in Giemsa-stained blood smears (*Fig. 6C; S6C,D*). This increase in pyknotic parasites occurred at the lowest levels of glycemia (*Fig. S6D*), suggesting that lowering glycemia, via the repression of HGP, induces *Plasmodium* pyknosis *in vivo*.

We than asked whether lowering glucose concentration is sufficient *per se* to induce parasite pyknosis. The percentage of human RBC infected *in vitro* by *P. falciparum* (*i*.*e*. parasitemia; % iRBC) was reduced in a dose dependent manner when glucose concentrations lower than 6 mmol/L (*i*.*e*. 110 mg/dL)(*Fig. 6E; S6E*). This was directly correlated with accumulation of pyknotic parasites (*Fig. 6F, Fig. S6E-G*) and by reduction of parasite mitochondrial function, revealed by loss of membrane potential (ΔΨm)(*Fig. 6G,H, Fig. S6H*). This suggests that repression of HGP arrests the development of asexual stages of *Plasmodium spp*. when blood glucose concentration falls below the 6 mmol/L (*i*.*e*. 110 mg/dL) threshold.

### Repression of HGP induces *P**lasmodium* sexual differentiation (*Fig. 6&S6*)

The number of circulating iRBC containing gametocytes was increased by 8.6-fold at the peak of *Pcc* infection in *G6pc1*^*Alb*Δ/Δ^ *vs. G6pc1*^*fl/fl*^ mice, as assessed by Giemsa-staining (*Fig. 6I, S6C-D*). This was corroborated by the accumulation of mRNA encoding gametocyte-specific genes, as assessed for *g377, gest* and *gep1* by qRT-PCR (*Fig. 6J*), confirming that repression of HGP induces gametocytogenesis and production of circulating gametocytes

### HGP sustains thermoregulation irrespectively of BAT thermogenesis (*Fig. 7*)

We asked whether repression of HGP controls the thermoregulatory response to *Pcc* infection via a mechanism involving brown adipose tissue (BAT) thermogenesis (Enerbäck et al., 1997). *Pcc* infection was associated with repression of BAT thermogenesis, to a similar extent in *G6pc1*^*Alb*Δ/Δ^ and *G6pc1*^*fl/fl*^ mice (*Fig. 7A,B*). However, this was not associated with a corresponding decrease of tail temperature in *Pcc*-infected *G6pc1*^*Alb*Δ/Δ^ mice (*Fig. 7B*), suggesting that heat dissipation (*i*.*e*. difference between tail and core body temperature) was higher in *Pcc*-infected *G6pc1*^*Alb*Δ/Δ^ *vs. G6pc1*^*fl/fl*^ mice (*Fig. 7C*).

**Figure 7.**
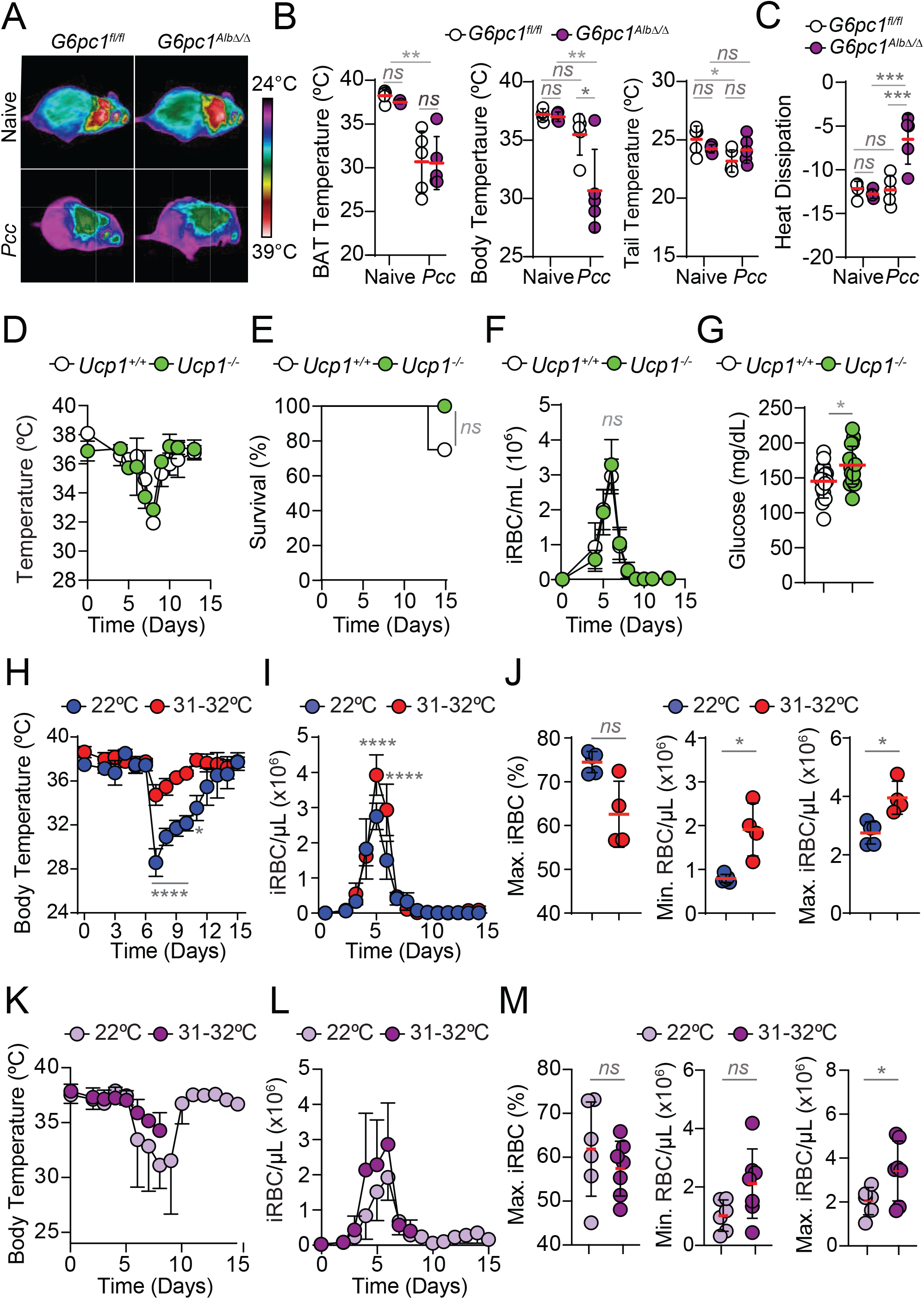
Glucose and temperature regulate *Plasmodium* behavior *in vivo*. (*see also Figure S7*) (A) Representative infrared images from N=4-5 mice *per* genotype, at steady state (naïve) and 7 days after *Pcc* infection. (B) Quantification of BAT, core body and tail temperatures in the same mice as (A). Data is represented as mean (red bars) ± SD, circles represent individual mice. (C) Heat dissipation (calculated as the difference between tail base and core body temperature) in the same mice as (A). Data is represented as mean (red bars) ± SD, circles represent individual mice. (D-G) Ucp1-dependent thermogenesis is dispensable for thermoregulatory response to *Pcc-*infection. (D) Core body temperature (mean ± SD) in *Ucp1*^*-/-*^ (N=4; green circles) *vs*. control *Ucp1*^*+/+*^ (N=4; white circles) mice. Data from one experiment. (E) Survival of *Pcc*-infected in the same mice as (D). (F) Pathogen load (mean ± SD) in the same mice as (D). (G) Blood glucose concentration, from the first 6 days of *Pcc* infection, in the same mice as (D). Data is represented as mean (red bars) ± SD, circles represent individual mice. (H-J) *Pcc*-infection in C57BL/6 mice housed at standard (22ºC; N=5; blue dots) or thermoneutral (31-32ºC; N=4; red dots) husbandry condition. (H) Core body temperature (mean ± SD). Data from one experiment. (I) Pathogen load (mean ± SD) in the same mice as (H). (J) Maximum parasitemia (% iRBC), minimum RBC numbers and maximum pathogen load (iRBC/µL), in the same mice as (H). Data is represented as mean (red bars) ± SD. Circles represent individual mice. (K-M) *Pcc*-infection in *G6pc1*^*Alb*Δ/Δ^ mice housed at standard (22ºC; N=5; blue dots) or at thermoneutral (31-32ºC; N=7; red dots) husbandry conditions. Data pooled from 2 independent experiments with the same trend. (K) Core body temperature represented as mean ± SD. (L) Pathogen load (mean ± SD) in the same mice as (K). (M) Maximum parasitemia (percentage iRBC), minimum RBC numbers and maximum pathogen load (iRBC/ µL), in the same mice as (K). Data is represented as mean (red bars) ± SD. Circles represent individual mice. P values in (B,C) determined using Two-Way ANOVA with Tukey’s multiple comparison, (D,F,H,I,K, L) determined using Two-Way ANOVA with Sidak’s multiple comparison, in (E) using Log-rank (Mantel-Cox) test (conservative) and in (G, J, M) using Mann Whitney test. *ns* – non-significant; * - p<0.05; ** - p<0.01; *** - p<0.001; **** - p<0.0001.

We than tested whether the thermoregulatory response to *Pcc* infection relies on the capacity of BAT to transform chemical energy into heat through a mechanism involving the mitochondrial uncoupling protein 1 (UCP1)(Enerbäck et al., 1997). *Ucp1* deletion (*Ucp1*^*-/-*^) did not affect the thermoregulatory response to *Pcc* infection (*Fig. 7D*) and had no impact on the survival (*Fig. 7E*) or pathogen load (*Fig. 7F*) in *Pcc*-infected *Ucp1*^*-/-*^ *vs. Ucp1*^*+/+*^ mice. However, reduction of glycemia was less pronounced in *Pcc*-infected *Ucp1*^*-/-*^ *vs. Ucp1*^*+/+*^ mice (*Fig. 7G*). This suggests that, in contrast to the thermoregulatory response to cold stress (Enerbäck et al., 1997) or to bacterial infection (Ganeshan et al., 2019), BAT thermogenesis is not essential to sustain core body temperature, while contributing to reduce glycemia during *Pcc* infection.

### Host glycemia and Core body temperature regulate *P**lasmodium* behavior *in vivo* (*Fig. 7&S7*)

To investigate further the mechanism via which glycemia regulates *Plasmodium* behavior, C57BL/6 mice were rendered hyperglycemic (*i*.*e*., glucose > 400 mg/dL) using streptozocin (STZ) (*Fig. S7A*). Glycemia was reduced in response to *Pcc* infection, to the same levels in STZ-treated *vs*. control mice (*Fig. S7A*). Moreover, hyperglycemia was restored upon *Pcc* clearance in STZ-treated mice (*Fig. S7A*), suggesting that insulin production by β-cells of the pancreas (i.e. destroyed by STZ) does not regulate glycemia in response to *Plasmodium* infection.

Hyperglycemic mice developed higher pathogen loads, compared to control mice (*Fig. S7B,C*). This was not associated with increased mortality (*Fig. S7D*) nor with changes in core body temperature (*Fig. S7E*), body-weight (*Fig. S7F*) or severity of anemia (*Fig. S7G*). This strengthens the idea that reducing glycemia, via repression of HGP, restrains the development of *Plasmodium* asexual stages.

To dissect the relative impact of glycemia *vs*. core body temperature on *Plasmodium* behavior, we considered that mice are not at thermoneutrality when maintained at standard husbandry conditions (22^0^C), allocating a significant proportion of EE to sustain core body temperature (Ganeshan and Chawla, 2017). At thermoneutral conditions (31-32^0^C), *Pcc* infected C57BL/6 mice reduced core body temperature to a lesser extent, compared to controls maintained at standard (22^0^C) conditions (*Fig. 7H*). This was associated with an increase in pathogen load (*Fig. 7I,J*) but not with increased mortality (*Fig. S7H*). Moreover, under thermoneutral conditions *Pcc*-infected mice did not reduce glycemia (*Fig. S7I*) or body weight (*Fig. S7J*) to the same extent as mice maintained under standard conditions and exhibited less severe anemia (*Fig. S7K*). This shows that re-allocation of EE from thermoregulation towards anabolic processes sustaining glycemia, RBC numbers and body weight occur at the cost of increased pathogen load. This suggests therefore that *Plasmodium* senses and reacts to the host’s core body temperature.

To examine the relative impact of glycemia and host core body temperature on the development of asexual stages of *Plasmodium*, the outcome of *Pcc* infection was compared in *G6pc1*^*Alb*Δ/Δ^ mice maintained at thermoneutral *vs*. standard conditions. At thermoneutrality, *Pcc*-infected *G6pc1*^*Alb*Δ/Δ^ mice decreased core body temperature to a lesser extent than *G6pc1*^*Alb*Δ/Δ^ mice housed at standard temperature (*Fig. 7K*). This was associated with an increase in pathogen load (*Fig. 7L,M*) and a borderline increase in mortality (*Fig. S7L*). Of note, *Pcc*-infected *G6pc1*^*Alb*Δ/Δ^ mice had similar glycemia (*Fig. S7M*), body weight loss (*Fig. S7N*) and severity of anemia (*Fig. S7O*) at thermoneutral *vs*. standard conditions. These observations suggest that repression of HGP induces a hypometabolic state whereby lowering body temperature contributes critically to modulate *Plasmodium* behavior.

## Discussion

*Pcc* infection in C57BL/6 mice is associated with the development of anorexia (*Fig. 1A,B*) and the reduction of blood glucose concentration (*Fig. 1C*)(Cumnock et al., 2018; Elased and Playfair, 1994). The latter relies on the repression of hepatic *G6pc1* expression (*Fig. 1D,E*) and HGP (*Fig. 1F*), which is restored over time to prevent the development of hypoglycemia (*Fig. 1I-L*) from compromising the energetic demands of the infected host (*Fig. 2A-H*). While perceived as a pathologic event *per se*, this hypometabolic state represents an evolutionary trade-off (Stearns and Medzhitov, 2015) of an otherwise adaptive response to *Plasmodium* spp. infection, whereby host and pathogens compete to access vital metabolites and micronutrients, such as iron and glucose, in detriment of each other.

Asexual stages of *Plasmodium spp*. can consume up to an estimated 60-80% of RBC Hb content, detoxifying excess heme and its associated iron in the form of hemozoin crystals (Sigala and Goldberg, 2014). The remaining Hb content is released upon hemolysis (Francis et al., 1997) as Hb (α2β2) tetramers, which disassemble into (αβ) dimers in plasma, releasing non-covalently bound heme (Hebbel et al., 1988; Pamplona et al., 2007) and producing labile heme (Ademolue et al., 2017; Elphinstone et al., 2016; Gouveia et al., 2017). As it accumulates in plasma, labile heme acts as an alarmin (Soares and Bozza, 2016), sensed by host pattern recognition receptors (Dutra and Bozza, 2014) and promoting the pathogenesis of severe forms of malaria (Knackstedt et al., 2019; Pamplona et al., 2007). When it accumulates in plasma at relatively low levels however, labile heme can elicit a protective response against severe malaria, such as the one underlying the beneficial effect of sickle Hb against malaria (Ferreira et al., 2008). Our current findings are consistent with this dual (*i*.*e*. pathologic *vs*. protective) effect of labile heme, whereby labile heme induces a hypometabolic state that restricts the development of asexual stages of *Plasmodium* while promoting malaria-associated hypoglycemia. In support of this idea, labile heme is sufficient to recapitulate the hallmarks of this hypometabolic state in non-infected mice, inducing a transient state of anorexia (*Fig. 1B and 2I,J*), repressing HGP (*Fig. 1D-F and 3F-H*), reducing glycemia (*Fig. 1C*)(Weis et al., 2017) and decreasing EE (*Fig. 2M,N*).

Labile heme induces this hypometabolic state via a mechanism that relies on the transcriptional repression of hepatic metabolic genes, including *G6pc1* (*Fig. 3C-H*). This occurs, most likely, via the inhibition one or several members of the SP/KLF family of transcription regulators (*Fig. S3G-I*), including SP1, which sustains *G6pc1* transcription (Wasner et al., 2001). SP1 can bind to the heme sensor Rev-erbα (Negoro et al., 2013), a transcriptional repressor (Raghuram et al., 2007) that inhibits *G6pc1* transcription (Yin et al., 2007). We speculate that upon heme sensing, Rev-erbα/β binds to SP1 and/or other SP/KLF family in the *G6pc1* promoter (Zhang et al., 2015), repressing *G6pc1* transcription and HGP.

Repression of HGP in response to *Pcc* infection is protective against liver damage (*Fig. 4H,I*), a hallmark of severe malaria (Knackstedt et al., 2019; Miller et al., 2002). The mechanism via which this occurs is most likely linked to a reduction in splenic NK and NKT cells and IFN-γ production (*Fig. 4A-G*). This is consistent with IFN-γ production by activated NK cells relying on glycolysis (Donnelly et al., 2014) and promoting organ damage during severe malaria (Franklin et al., 2009; Su and Stevenson, 2000).

This immunosuppressive effect did allow *Pcc* immune evasion (*Fig. 1K,L*) and was associated instead with marked repression of the development of asexual stages of *Pcc* (*Fig. 5A,B*). This suggests that the hypometabolic state imposed by labile heme, via the repression of HGP and the reduction of glycemia, is a component of innate nutritional immunity (Nunez et al., 2018). Presumably, this defense strategy restrains asexual stages of *Plasmodium spp*. from accessing not only glucose, but perhaps other essential host-derived metabolites and micronutrients, such as amino acids (*e*.*g*. isoleucine or glutamine) or lipids (*i*.*e*. lysophosphatidylcholine)(Kumar et al., 2021; Mancio-Silva et al., 2017). This interpretation is consistent with glycolysis being required to support the high energetic demands and anabolic metabolism inherent to the proliferation of asexual stages of *P. falciparum* (Homewood, 1977; Meireles et al., 2017; Olszewski and Llinas, 2011). Moreover, glycolysis is also essential to fuel *P. falciparum* pentose phosphate pathway and maintain parasite redox balance and viability (Preuss et al., 2012a).

Repression of HGP is sensed, directly or indirectly, by asexual stages of *Plasmodium* and reacted upon via a transcriptional response associated with the induction of parasite pyknosis (*Fig. 6C,D, S6C,D*). This is paralleled by the induction of sexual differentiation, characterized by the generation of high numbers of circulating gametocytes (*Fig. 5E-G, 6I,J*), a reliable predictor of malaria transmission (Mackinnon and Read, 1999). This suggests that repression of HGP and hypoglycemia favor malaria transmission.

The observation that repression of HGP induces *Plasmodium* sexual commitment, presumably to favor malaria transmission (*Fig. 5E-G, 6I,J*), while reducing virulence (*Fig. 5H, 6A,B*) is reminiscent of that observed during bacterial infection, whereby iron reprograms host glucose metabolism to regulate bacterial transmission *vs*. virulence (Rao et al., 2017; Weis et al., 2017). This suggests that cross-regulation of host iron and glucose metabolism, is an evolutionarily conserved defense strategy against infection by different classes of pathogens.

Our interpretation is that reprogramming of host iron-heme metabolism in response to infection induces a coordinated response, whereby anorexia is coupled to the repression of HGP to reduce glycemia and deprive asexual stages of *Plasmodium* from accessing glucose, impairing parasite development and inducing parasite pyknosis (*Fig. 6C-F, S6C-G*). This defense strategy carries two major trade-offs, namely the development of malaria-associated hypoglycemia, an independent risk factor of malaria mortality (Madrid et al., 2015; Marsh et al., 1995; Service, 1995; White et al., 1983) and a possible increase in parasite transmission.

HGP regulates the thermoregulatory response to malaria (*Fig. 7A-C*), possibly interfering with the development of asexual stages of *Plasmodium spp*. (*Fig. 7H-M*)(Rojas and Wasserman, 1993). In contrast to humans, which increase body temperature, *i*.*e*. fever, as a cardinal sign of *Plasmodium spp*. infection (Evans et al., 2015), mice appear to suppress HGP and reduce core body temperature as an alternative strategy to restrain the development and proliferation of asexual stages of *Plasmodium* (*Fig. 7H-M*), in keeping with thermoregulation being an evolutionarily conserved defense strategy against infection (Schieber and Ayres, 2016).

In contrast to the canonical thermoregulatory response to cold stress (Enerbäck et al., 1997) or to systemic bacterial infection (Ganeshan et al., 2019), the thermoregulatory response to *Pcc* infection does not rely on UCP1-dependent BAT thermogenesis (*Fig. 7A-G*). It occurs instead via a mechanism whereby repression of HGP limits glucose supply to support heat generation inherent to organismal metabolism (*Fig. 2A,B*) while sustaining heat dissipation (*Fig. 7A-C*). This strategy carries a major risk of core body temperature falling below a threshold compatible with host survival. This is likely aggravated by voracious glucose consumption by asexual stages of *Plasmodium* (Olszewski and Llinas, 2011; Thien et al., 2006) as well as by proliferating immune cells (Buck et al., 2017; Man et al., 2017; Wang et al., 2019).

In conclusion, labile heme orchestrates a host hypometabolic response that restricts asexual stages of *Plasmodium* from accessing glucose, a vital nutrient supporting parasite development and proliferation. When sustained over time this host defense strategy becomes a driving force in the pathogenesis of malaria-associated hypoglycemia. Asexual stages of *Plasmodium* can sense the development of hypoglycemia either directly, or indirectly via the associated decrease of core body temperature, activating a transcriptional program that reduces virulence and promotes sexual differentiation, presumably as a strategy favoring malaria transmission.

## ACKNOWLEDGEMENTS

We are indebted to Dr. Silvia Portugal (Max Plank Institute, Berlin) and Dr. Jessica Ann Thompson (Instituto Gulbenkian de Ciência) for critical review of the manuscript, Dr. Joanne Thompson (University of Edinburgh) for the PcAS-GFP_ML_ parasites; BEI Resources for *P*. falciparum parasites (MRA-1029, deposited by Andrew M. Talman and Robert E. Sinden), excellent support from IGC’s Advanced Imaging; Antibody; Flow Cytometry and Genomics facilities. SR was supported by Fundação para a Ciência e Tecnologia (FCT; 5723/2014; FEDER029411), TWA by Gulbenkian foundation (IBB2017), EJ by the Deutsche Forschungsgemeinschaft (DFG, EXC 2051; 390713860), ARC by FCT (SFRH/BPD/101608/2014), SW and JG by the by the Center for Sepsis Control and Care (CSCC), Jena University Hospital (BMBF 01EO1502) and DFG (EXC 2051; 390713860 and WE 4971/6-1), DD and FN were supported by FCT through GHTM (UID/04413/2020). The MPS laboratory at Instituto Gulbenkian de Ciência is supported by the Gulbenkian Foundation and by “La Caixa” (HR18-00502) and FCT (5723/2014; FEDER029411) foundations. MPS is an associate member of the DFG Cluster of Excellence ‘Balance of the Microverse’ (https://microverse-cluster.de/en). Support by Congento (LISBOA-01-0145-FEDER-022170) is acknowledged.

## AUTHOR CONTRIBUTION

SR and TWA contributed critically to study design, experimental work, data analyzes and interpretation. EJ designed (with TWA), performed *Plasmodium* scRNA and analyzed data (with SR, AS and JL). QW carried *in vitro* experiments on heme regulation of gluconeogenesis, following initial experiments by ARC. JG and SW performed indirect calorimetry measurements of heme-treated mice and analyzed data. RM designed and performed flow cytometry experiments (with TWA). GP assisted SR and TWA in experiments related to *Plasmodium* virulence. IM and ES assisted SR and TWA in experiments related to the induction of anorexia in response to *Plasmodium* infection. SC maintained and characterized mouse strains. TWA performed bulk-RNA sequencing experiments and contributed to analyzes by AS and JL. GM and FR participated in data interpretation. FN designed, performed and analyzed *P. falciparum* glucose tolerance *in vitro* experiments (with SR). DD assisted FN in *in vitro* experiments. MPS formulated the original hypothesis, designed the study and wrote the manuscript with SR and TWA. All authors read and approved the manuscript.

## SUPPLEMENTARY FIGURE LEGENDS

**Figure S1. HGP controls glycemia in response to *Pcc* infection**. (*Related to Figure 1*).

(A) Pyruvate tolerance test (PTT) in control (Ctrl.; fasted overnight; N=4) and *Pcc*-infected (*i*.*e*. peak of infection, day 7; N=5) C57BL6 mice, at the times indicated after pyruvate administration (*left panel*) and corresponding area under the curve (*right panel*). Data represented as mean ± SD, from one experiment.

(B) Incidence of seizures after insulin administration in control and *Pcc*-infected C57BL6 mice. Same mice as *Fig. 1F*.

(C) Blood glucose concentration, represented as mean ± SD, in *Pcc*-infected *G6pc1*^*Alb*Δ/Δ^ (N=15; purple circles) and *G6pC1*^*fl/lfl*^ (N=12; white circles) mice. Same mice as *Fig. 1G-J*. Gray values are the proportion of surviving over total number of mice, at indicated time points.

(D) Insulin concentration in plasma of *G6pc1*^*Alb*Δ/Δ^ (N=5; purple circles) and *G6pC1*^*fl/lfl*^ (N=5; white circles) mice at the peak of *Pcc* infection (*i*.*e*. day 7). Data represented as mean ± SD from one experiment. Gray area: physiologic range.

(E-J) Disease parameters in same mice as *Fig. 1G-J*.

(E) Core body temperature, represented as mean ± SD.

(F) Disease trajectories established by the relationship between core body temperature and parasite density (iRBC/µL). Circles represent death.

(G) Minimum core body temperature throughout *Pcc* infection. Data represented as mean (red bars) ± SEM. Circles represent individual mice.

(H) Percentage of initial body weight. Data is represented as mean ± SEM. Gray values are the proportion of surviving over total number of mice, at indicated time points.

(I) Disease trajectories established by the relationship between percentage of initial body weight and number of iRBC per µL. Circles represent death.

(J) Minimum percentage of initial body weight throughout *Pcc* infection. Data represented as mean (red bars) ± SEM. Circles represent individual mice.

*P* values determined in: (A) using Two-Way ANOVA with Tukey’s multiple comparison test for Glucose concentration and, t test for AUC, in (B) using Log-rank

(Mantel-Cox) test (conservative), in (D,G,J) using Mann-Whitney U test, (E,H) using Two-Way ANOVA with Sidak’s multiple comparison test. *ns* – non-significant; *p<0.05; **p<0.01; ***p<0.001.

**Figure S2. HGP controls host EE during *Plasmodium* infection**. (*Related to Figure 2*)

(A) Average of fine movements (*i*.*e*. non-directed ambulatory motion, expressed as: All distances traveled (i.e. Total Activity) - Directed ambulatory locomotion)) in the period highlighted by light blue in (*Fig. 2A*), segregated into one light/dark cycle. Data represented as mean ± SEM, from the same mice as *Fig. 2A-H*.

(B-H) Metabolic and behavioral parameters of naïve *G6pc1*^*Alb*Δ/Δ^ (N=4; purple circles) and *G6pC1*^*fl/lfl*^ (N=4; white circles) mice, represented as mean ± SEM, pooled from 2 independent experiments.

(B) Energy expenditure (EE) and (C) mean hourly EE, in the 3 day-period represented in (B), segregated into daily light/dark cycle.

(D) Average of food intake in the 3 days time-period represented in (B), segregated into daily light/dark cycle.

(E) Respiratory exchange ratio (RER) and (F) mean RER, in the 3 day-period represented in (E), segregated into daily light/dark cycle.

(G) Means VCO_2_ in the 3 day-period represented in (E), segregated into daily light/dark cycle.

(H) Means VO_2_ in the 3 day-period represented in (E), segregated into daily light/dark cycle.

(I-N) Metabolic and behavioral parameters of C57BL/6 mice, receiving heme (N=4; brown circles) or control PPIX (N=4; white circles) at the time indicated by the arrows, from the same mice as *Fig. 2I-P*.

(I) Fine movement (*i*.*e*. non-directed ambulatory motion) and (J) Means of fine movements in the 24h period highlighted by light blue in (I), segregated into one light/dark cycle.

(K) VO_2_ and (L) Means of VO_2_, corresponding to the 24h period highlighted by light blue shading in (K), segregated into one light/dark cycle.

(M) VCO_2_ and (N) average VCO_2_ in the 24h period highlighted by light blue shading in (M), segregated into one light/dark cycle.

(O) Glucose concentration after pyruvate administration (2mg/g of BW) test in C57BL6 mice receiving heme (N=4) or vehicle controls (N=5). Mean glycemia represented as mean ± SD, from one experiment.

*P* values in (N) were determined using student t-test and in the remaining panels using Two-Way ANOVA with Sidak’s multiple comparison test. *ns* – non-significant; * p<0.05; **p<0.01; ****p<0.0001.

**Figure S3. Labile heme recapitulates the repression of HGP in response to *Plasmodium* infection**. (*Related to Figure 3*)

(A) Principal component analysis (PCA) of 18,136 transcripts differentially expressed in whole livers from C57BL/6 mice subjected to fasting (fasted overnight; 15h; N=4), receiving heme (30 mg/Kg, overnight; N=3) or at the peak of *Pcc* infection (*i*.*e*. day 7; N=4).

(B) Scatter plot of the 18,136 transcripts differentially expressed in the liver of the same mice as (A). Plotting log2 fold change (FC) of *Pcc* infection *vs*. fasting (x-axis) against log2FC of heme administration *vs*. fasting (y-axis), was used to highlight shared transcripts (adjusted p-value <0.05). Upper right quadrant contains shared up-regulated transcripts (1551), lower left quadrant contains shared downregulated transcripts (1747). Shared transcripts had an absolute log2 fold change higher than 1, compared to fasted controls, in both pairwise comparisons. Broken red lines marks LogFC = 1.

(C) Venn diagram of pathways (*i*.*e*. GO terms database) downregulated by heme administration and by *Pcc*-infection, as compared to fasted control mice.

(D) Functional enrichment analysis of the 12 up-regulated pathways (*i*.*e*. GO terms database) with highest enrichment scores (−log10 adjusted p-value) in *Pcc* infected *vs*. control fasted mice.

(E) Functional enrichment analysis of the 12 up-regulated pathways (*i*.*e*. GO terms database) with highest enrichment score (−log10 adjusted p-value) in mice receiving heme *vs*. control fasted mice.

(F) Venn diagram of pathways, using GO terms database, up-regulated by heme administration and by *Pcc*-infection, as compared to control fasted mice.

(G) Functional enrichment analysis of DNA-binding motifs (*i*.*e*. TRANSFAC database) in the regulatory regions of shared transcripts downregulated by heme administration and *Pcc*-infection, compared with control fasted mice respectively. Left panel illustrates DNA-binding motifs with the highest enrichment score (−log10 adjusted p-value)(*left panel*). Right panel illustrates DNA-binding motifs with the highest enrichment score (−log10 adjusted p-value) downregulated exclusively in response to heme administration or to *Pcc*-infection.

(H) Venn diagram representing the number and percentage of DNA-binding sites present in the regulatory regions of shared transcripts downregulated by heme administration and *Pcc*-infection, compared with fasted control mice, respectively.

(I) Transcription factor motif prediction analysis (*i*.*e*. JASPAR) of mouse *G6pc1* promoter (−320 to +20), highlighting DNA-binding sites also present in the regulatory regions of shared transcripts downregulated by heme administration and *Pcc*-infection, compared with fasted control mice respectively (as in G).

**Figure S4. Repression of HGP exerts no effect on monocyte/macrophages in response to *Pcc* infection**. (*Related to Figure 4*)

(A-D) Analyzes of splenic monocyte/macrophages from naïve (N=4 per genotype) and *Pcc*-infected (*i*.*e*. peak of infection, day 7; N=6 per genotype) *G6pc1*^*Alb*Δ/Δ^ (purple dots) and *G6pC1*^*fl/lfl*^ (white dots) mice. Data is represented as mean (red bar) ± SD from 2 independent experiments with similar trend. Circles represent individual mice. Same mice as in *Fig. S4A-F*.

(A) Representative FACS plots for splenic CD11b and F4/80 staining (*i*.*e*. gated on (CD19^-^Ly6G^-^).

(B) Quantification of splenic monocyte/conventional dendritic cells (CD11b^+^F4/80^-^).

(C) Quantification of splenic monocytes (CD11b^+^F4/80^int^).

(D) Quantification of splenic red pulp macrophages (CD11b^-^F4/80^+^).

(E) Quantification of circulating neutrophils in *G6pc1*^*Alb*Δ/Δ^ (purple dots; N=5) and *G6pC1*^*fl/lfl*^ (white dots; N=5) mice, at the peak of *Pcc*-infection. Same mice as in *Fig. 5C*. Data is represented as mean (red bar) ± SD from one experiment. Circles represent individual mice.

(F) Cytokine/chemokine concentration (pg/mL) in the plasma of naïve (N=5 per genotype) and at the peak (*i*.*e*. day 7) of *Pcc* infection in *G6pc1*^*Alb*Δ/Δ^ (N=12; purple circles) and *G6pC1*^*fl/lfl*^ (N=11; white circles) mice. Same mice as in *Fig. 4G*. Data is represented as mean ± SD and from 2 independent experiments with similar trend.

*P* values in (B-D, F) were determined using Two-Way ANOVA with Tukey’s multiple comparison test. In E using unpaired t test. *ns* – non-significant; * - p<0.05.

**Figure S5. Repression of HGP regulates *Plasmodium* behavior**. (*Related to Figure 5*)

(A) Hemograms at the peak of *Pcc* infection (*i*.*e*. day 7) in *G6pc1*^*Alb*Δ/Δ^ (N=5; purple) and *G6pC1*^*fl/lfl*^ (N=5; white) mice (same mice as in *Fig. 5C*), depicting erythrocyte numbers, hemoglobin concentration, hematocrit, RBC mean corpuscular volume (MCV), RBC mean corpuscular hemoglobin (MCH), RBC mean corpuscular hemoglobin concentration (MCHC) and RBC distribution width (RDW). Data represented as mean (red bar) ± SD from one experiment. Circles represent individual mouse. Gray rectangles indicate range of reference values for steady state. P values were determined using unpaired t test. *ns* – non-significant; * - p<0.05, ** - p<0.01.

(B) Expression pattern of parasite stage-specific transcripts across the combined single-cell (sc) RNA UMAP projection of transcriptomes in FACS-sorted iRBCs from *G6pc1*^*fl/lfl*^ and *G6pc1*^*Alb*Δ/Δ^ mice. In ring stages: enolase (*eno*, PCHAS_1215000) and succinate dehydrogenase subunit 4 (*sdh4*, PCHAS_1209400); in trophozoites: phosphoglycerate kinase (*pgk*, PCHAS_0823700) and *fam a* (PCHAS_120120); in schizonts: rhoptry neck protein 2 (*ron2*, PCHAS_1319000) and merozoite surface protein 9 (*msp9*, PCHAS_1445500), in sexually committed (SC) rings: carbon catabolite repression-negative on TATA-less, (CCR4-Not) subunit 2 (*not2*, PCHAS_0924800) and in gametocytes: SOC protein 3 (*soc3*, PCHAS_1307700).

(C) Combined UMAP projection of mouse single RBC transcriptomes in the same FACS-sorted iRBCs from *G6pc1*^*fl/lfl*^ and *G6pc1*^*Alb*Δ/Δ^ mice in (B). Each dot represents a transcriptome from one individual RBC.

(D) Functional enrichment analysis of parasite pathways (*i*.*e*. GO terms database) down regulated in Cluster 3 (*i*.*e*. sexually-committed rings) in FACS-sorted iRBCs from *G6pc1*^*Alb*Δ/Δ^ *vs. G6pc1*^*fl/lfl*^ mice.

(E) Functional enrichment analysis of parasite pathways (*i*.*e*. GO terms database) upregulated in Cluster 4 (*i*.*e*. trophozoites) in FACS-sorted iRBCs from *G6pc1*^*Alb*Δ/Δ^ *vs. G6pc1*^*fl/lfl*^ mice.

(F) Functional enrichment analysis of parasite pathways (*i*.*e*. GO terms database) upregulated in Cluster 7 (*i*.*e*. rings) in FACS-sorted iRBCs from *G6pc1*^*Alb*Δ/Δ^ *vs. G6pc1*^*fl/lfl*^ mice.

**Figure S6. Repression of HGP reduces *Plasmodium* virulence and induces gametocytogenesis**. (*Related to Figure 6*)

(A, B) Repression of HGP lowers *Pcc* virulence. *G6pc1*^*Alb*Δ/Δ^ mice were infected with *Pcc* iRBC from *G6pc1*^*Alb*Δ/Δ^ (purple circles) *vs. G6pc1*^*fl/lfl*^ (light purple circles) mice (N=6 per group) (same mice as *Fig. 6B*). Data pooled from 2 independent experiments with similar trend.

(A) Body temperature (*left panel*) and minimum body temperature throughout the 15-days of *Pcc* infection (*right panel*) represented as mean ± SD. Circles in right panel represent individual mice and red bars represent mean.

(B) Glycemia (*left panel*) and minimum glycemia throughout the 15 days of *Pcc* infection (*right panel*) presented as mean ± SD. Circles in right panel represent individual mice and red bars in right panel mean.

(C, D) Representative Giemsa-stained thin blood smears of *G6pc1*^*fl/lfl*^ (C; N=14) and (C) (C) (C) *G6pc1*^*Alb*Δ/Δ^ (D; N=17) mice, 7 days after *Pcc* infection. Data representative of 3 independent experiments, with similar trend. Parasite stages were identified based on morphology. Arrows highlight parasites, purple text: blood glucose concentration. Scale bar is 2µm.

(E-H) Glucose concentration regulates *P. falciparum* development and proliferation *in vitro*.

(E) Representative Giemsa-stained thin blood smears of *P. falciparum* infected human RBC (iRBC) cultured *in vitro* under varying glucose concentrations *vs*. control media (∼11mmol/L Glucose). Indented square (0h) represents original seeded culture. Filled arrowheads indicate healthy parasites, empty arrowheads aberrant parasites and asterisks pyknotic parasites. Images representative of 3-4 replicates in 3 independent experiments. Scale bar is 2µm.

(F, G) Percentage of *P. falciparum* iRBC cultured *in vitro* for 24h (F) and 72h (G), under varying glucose concentrations (23-0 mmol/L), containing morphologically healthy, aberrant or pyknotic parasites. Data is from the same experiment as in (E) and presented as mean ± SEM, from 3 wells in 1 out of 3 independent experiments with similar trend.

(H) Gating strategy used to identify *P. falciparum* infected human RBC (iRBC). Cells were gated based on size and granularity (FCS-A *vs*. SSC-A plot). Single cells were identified based on FSC-A *vs*. FSC-H profile, and GFP or Hoechst (DNA content) signals were used to identify *P. falciparum* iRBC.

*P* values in (F,G) and for core body temperature and blood glucose in (A, B) were determined using Two-Way ANOVA with Tukey’s multiple comparison test and in (A,B) for minimum core body temperature and blood glucose using unpaired t test. *ns*

– non-significant; * - p<0.05; ** - p<0.01; *** - p<0.001; **** - p<0.0001.

Figure S7. Host glycemia and core body temperature regulate *Plasmodium* behavior *in vivo*. (*Related to Figure 7*)

(A-G) Hyperglycemia promotes *Pcc* expansion. hyperglycemia was induced by the administration of streptozocin (STZ) prior to *Pcc* infection in C57BL/6 mice STZ (N=3; purple dots). Controls (Ctr.) received vehicle (PBS; N=5; lilac dots). Red bars in (C) indicate mean. Data from one experiment.

(A) Blood glucose concentration presented as mean ± SD.

(B) Pathogen load presented as mean ± SD.

(C) Maximum percentage of iRBC, minimum RBC count and maximum count of iRBC. Data represented as mean (red bars) ± SD. Circles correspond to individual mice.

(D) Survival.

(E) Individual relationships (*i*.*e*. disease trajectories) between body temperature and pathogen load during the entire course of *Pcc* infection.

(F) Individual relationships (*i*.*e*. disease trajectories) between percentage of initial body weight and pathogen load during the entire course of *Pcc* infection.

(G) Individual relationships (*i*.*e*. disease trajectories) between the number of circulating RBC and pathogen load during the entire course of *Pcc* infection.

(H-K) Core body temperature regulates *Pcc* expansion. *Pcc*-infected C57BL/6 mice, housed at standard husbandry (22ºC; blue circles; N=5) or at thermoneutral condition (31-32ºC; red circles; N=4). Circles in (I-K) indicate death. Data from one experiment. Same mice as *Fig. 7-H-J*.

(H) Survival.

(I) Individual relationships (*i*.*e*. disease trajectories) between Glycemia and pathogen load during the entire course of *Pcc* infection.

(J) Individual relationships (*i*.*e*. disease trajectories) between percentage of initial body weight (compared to non-infected) and pathogen load during the entire course of *Pcc* infection.

(K) Individual relationships (*i*.*e*. disease trajectories) between the number of circulating RBC and pathogen load during the entire course of *Pcc* infection.

(L-O) Glycemia and core body temperature regulates *Pcc* expansion. *Pcc*-infected of *Pcc*-infected *G6pc1*^*Alb*Δ/Δ^ mice housed at standard husbandry (22ºC; violet circles; N=5) or at thermoneutral condition (31-32ºC; purple circles; N=7). Circles in (M-O) indicate death. Data pooled from 2 independent experiments. Same mice as *Fig. 7-K-M*.

(L) Survival.

(M) Individual relationships (*i*.*e*. disease trajectories) between Glycemia and pathogen load during the entire course of *Pcc* infection.

(N) Individual relationships (*i*.*e*. disease trajectories) between percentage of initial body weight and pathogen load during the entire course of *Pcc* infection.

(O) Individual relationships (*i*.*e*. disease trajectories) between the number of circulating RBC and pathogen load during the entire course of *Pcc* infection.

P values determined in (A,B) using Two-Way ANOVA with Sidak’s multiple comparison test, in (C) using Mann Whitney test, in (D,H,L) using Log-rank (Mantel-Cox) test (conservative). *ns* – non-significant; *p<0.05; ** - p<0.01; ***p<0.001; ***p<0.0001.

## STAR METHODS

### Key Resources Table (see Key Resources file)

#### Contact for Reagent and Resource Sharing

Further information and requests for resources and reagents should be directed to and will be fulfilled by the Lead Contact, Miguel P. Soares.

#### Experimental Model and Subject Details

##### Mice and Conditional allele deletions

Mice were bred and maintained under specific pathogen-free (SPF) conditions at the Instituto Gulbenkian de Ciência (IGC) or at the Jena University Hospital. Mice were housed at standard vivarium temperature (22°C) or at thermoneutrality (31-32°C) in a 12-hour light/dark cycle with free access to water and standard chow pellets. Experimental protocols were approved by the Instituto Gulbenkian de Ciência Ethics Committee/ORBEA and the Portuguese National Entity (Direcção Geral de Alimentação e Veterinária) or by the Jena University Hospital regional animal welfare committee (Registration number: 02-048/16, Thuringian State Office for Consumer Protection and Food Safety). Experimental procedures were performed according to the Portuguese or the German legislation on protection of animals and European (Directive 2010/63/EU) legislations. C57BL/6J mice were obtained from the IGC or the Jena University Hospital animal facilities. C57BL/6 *Sa*^CreERT2/Wt^*G6pc1*^fl/fl^ (i.e. *G6pc1*^*Albfl/fl*^) mice and littermate control *G6pc1*^*fl/fl*^ mice were previously described (Mutel et al., 2011; Weis et al., 2017). Conditional deletion of the *G6pc1*^fl/fl^ allele in hepatocytes was induced by Tamoxifen administration (*i*.*p*., 10 mg/mL, 100μL; Sigma), daily for 5 days, starting at 8-week after birth as described (Mutel et al., 2011; Weis et al., 2017). The resulting *G6pc1*^*Alb*Δ/Δ^ mice were used 2 weeks after deletion. *Ucp1*^*-/-*^ (B6.129-Ucp1^tm1Kz^/J) mice were obtained from The Jackson Laboratory (JAX #003124) and breed in heterozygosity at the IGC animal house. Wild type littermates were used as controls.

##### Plasmodium

Mice were infected with the following *Plasmodium chabaudi chabaudi* (*Pcc*) strains: *Pc*AS (Reece and Thompson, 2008) and transgenic GFP-expressing *Pc*AS (PcAS-GFP_ML_)(Marr et al., 2020; Ramos et al., 2019). *In vitro* assays were performed using a transgenic GFP-expressing, laboratory-adapted *P. falciparum* line 3D7-GFP (*Pf*-3D7-GFP; MRA-1029, MR4, ATCC® Manassas Virginia).

#### Method details

##### Plasmodium chabaudi chabaudi infection and disease assessment

Infections were performed as described (Ramos et al., 2019). Briefly, blood was harvested from an infected mouse carrying ∼10%, parasitemia, diluted in PBS to reach ∼1×10^7^ iRBC/mL and an inoculum of 200 µL (∼2×10^6^ iRBC) was administered (i.p.). Disease progression was monitored daily, from day zero (D0) onwards quantifying anorexia, body weight (Ohaus^®^ CS200 scaler, Sigma Aldrich), core (*i*.*e*. rectal) body temperature (Rodent thermometer; BIO-TK8851, Bioset) and blood glucose concentration (Accu-CHECK Performa glucometer, Roche), as well as parasitemia and number of RBC, as described (Ramos et al., 2019).

##### Food intake

Steady-state daily food intake was quantified for 2 days before infection, and subsequently throughout the course of infection. Briefly C57BL/6 mice were housed in individual cages with free access to water and standard chow pellets. Food pellets were weighed (Ohaus^®^ CS200 scaler, Sigma Aldrich) daily at 6:00pm and daily food consumption was calculated as follows: Pellet Mass_24hrs before_ - Pellet Mass _24hrs after_ = daily food intake.

##### Infrared Temperature Measurements

BAT and tail temperatures were measured from digital images obtained using a infrared camera (FLIR E96: Compact-Infrared-Thermal-Imaging-Camera; FLIR Systems; West Malling, Kent, UK), analyzed with a software package (FLIR Tools^®^ Software, FLIR Systems; West Malling, Kent, UK) as described (Blankenhaus et al., 2019). Before acquiring images, mice were anesthetized (1-2% Isoflurane) and the interscapular area was shaved, after which mice were allowed to recover for at least 24 hours. Several infrared images of the BAT and Tail base were taken while mice were allowed to move freely on a cage lid. The analysis software was used to determine the average temperature of a selected region of the interscapular area and tail base across 3-4 images from the same animal. Images were acquired in non-infected, or *Pcc* infected (Day 7).

##### Pathogen load

Number of RBC per µL of mouse blood was quantified by flow cytometry (FACSCalibur analyzer; BD Biosciences) using a standard concentration of reference latex beads (10μm; Coulter® CC Size Standard L10, Beckman Coulter). Gating on RBC was based on size *vs*. granularity and on bead population. Percentage of *Pcc* AS iRBCs (*i*.*e*., parasitemia) was determined manually by optical microscopy, counting RBC in 4 fields of Giemsa-stained blood smears (1000x magnification). Percentage of PcAS-GFP_M_-iRBC was determined by flow cytometry, according to the percentage of RBC expressing GFP (GFP^+^). Pathogen load was determined as: Parasitemia x RBC number and is expressed as iRBC/µL. Disease trajectories were plotted using disease parameters *vs*. parasite density, daily for individual mice, over the entire period of the infection, as described (Ramos et al., 2019).

##### *Plasmodium falciparum in vitro* culture

Laboratory-adapted *Plasmodium* falciparum (*Pf*)-3D7-GFP was co-cultured (37°C; 95% humidity, 5% of CO_2_) with human erythrocytes from healthy donors (5% hematocrit), replacing human serum was replaced by 0.5% AlbuMAXII (Invitrogen™, Thermo Fisher Scientific), as described (Santos et al., 2015). Briefly, *Pf*-3D7-GFP cultures were synchronized using 5% D-Sorbitol (Sigma-Aldrich), returned to standard culture conditions until parasite RBC reinvasion, synchronized twice again, with a 6h interval and returned to culture (Lobo et al., 2018). Parasites at the schizont stage were suspended, layered on a 70% Percoll (Sigma-Aldrich), centrifuged (1000g; 15 min. no brake), collected from the upper layer of Percoll cell suspension, washed (PBS) and incubated in standard culture conditions until RBC reinvasion (Santos et al., 2015). Ring stage parasites (<10h post-invasion) were diluted to _≅_0,8% parasitemia and 3% hematocrit in RPMI 1640 without glucose (Gibco™, Thermo Fisher Scientific) and seeded on a 24-well plate. Parasites were then cultured under increasing D-glucose concentrations (Sigma-Aldrich, Thermo Fisher Scientific). Slides were prepared at 24, 48 and 72h and stained with Giemsa. The remaining of the parasites in the same culture were washed (PBS/BSA) and analyzed by flow cytometry (described below).

##### *Plasmodium* blood stages assessment

*Pcc* ring, trophozoite, schizont and gametocyte stages were quantified in Giemsa-stained blood smears by morphological assessment, based on DNA content, parasite size and hemozoin deposition. Pyknotic parasites were identified by their condensed chromatin mass without discernable cellular structures (*i*.*e*. cytoplasm, food vacuole) and hemozoin. The percentage of each *Plasmodium* stage was calculated over total number of iRBC and density was calculated as described above.

In *P. falciparum* assays, aberrant *Pf*-3D7-GFP parasites presented morphologically a compromised cell structure or delayed development while pyknotic *Pf*-3D7-GFP had condensed chromatin without discernable cytoplasm, food vacuole or hemozoin. These were compared to healthy parasites grown in a control media (RPMI1640, Biowest, routinely used for *P. falciparum* growth, glucose ∼11mmol/L concentration). Blood smear images were acquired on a Zeiss Imager Z2/ApoTome.2 system, equipped with an Axiocam 105 color camera, using the 100x 1.4NA Oil immersion objective, in a 5×5 tile stitched with Zeiss’s ZEN v3.1 and analyzed using the Fiji Software (ImageJ). Background in parasite RGB images was corrected using the Colour Correct plugin for Fiji, contributed by Gabriel Landini (https://github.com/landinig/IJ-Colour_Correct/blob/main/colour_correct.zip).

##### Plasmodium virulence assay

*Pcc* AS parasites were adoptively transferred from *G6pc1*^*Alb*Δ/Δ^ vs. *G6pc1*^*fl/fl*^ mice into *G6pc1*^*Alb*Δ/Δ^ or C57BL6/J mice. Briefly, *Pcc* infection was monitored at the peak of infection (D6-D7) to determine blood glucose concentration and the presence of pyknotic parasites in Giemsa-stained blood smears. At the time point when *G6pc1*^*Alb*Δ/Δ^ mice presented severe hypoglycemia and the highest number of RBC containing pyknotic parasites, blood was collected from both *G6pc1*^*Alb*Δ/Δ^ and *G6pc1*^*fl/fl*^ control mice. The blood of each donor animal was diluted so that an estimated 1.2-1.5×10^6^ iRBC were passively transferred (i.p.) into recipient *G6pc1*^*Alb*Δ/Δ^ or C57BL6/J mice. These were monitored daily for survival and disease assessment, as described above.

##### Glucose, Pyruvate and Insulin Tolerance Tests

Glucose (GTT; Sigma; 1 mg/g BW), pyruvate (PTT; sodium Pyruvate, Sigma; 2mg/g BW) and insulin (ITT; Humulin R U-100 UI/mL; 2 Units per mouse, Eli Lilly) tolerance tests were performed on *Pcc* infected (at day 7) *vs*. control overnight fasted C57BL6 male mice or on overnight-fasted C57BL/6 mice subsequently receiving heme (i.p., 30 mg/Kg BW) or vehicle (PBS) and tested after 10 hours. Blood was drawn from a tail cut and blood glucose concentration was measured using a glucometer (AccuCheck System, Roche) at 0, 15, 30, 60, 90, and 120 mins after injection.

##### Heme administration

Hemin (Frontier Scientific) stock solutions were prepared, as described (Ramos et al., 2019). Briefly, hemin was dissolved in 0.2N NaOH and buffered to pH 7.4 using 0.2N HCl. After filtration (70µm cell strainer) to remove precipitates, a stock solution was stored at -80ºC up to one month. Heme absorbance was determined using a spectrophotometer (SmartSpec 3000) at λ_405nm_ and concentration was calculated using extinction coefficient (E_mM_ = 85.82), following the Lambert-Beer law (A_405nm_=εcl). Heme was diluted in PBS and administered (i.p., 30 mg/Kg BW).

##### Promethion behavioral and phenotyping system

Promethion Core (Sable Systems, USA) was used to measure indirect calorimetry. Mice were housed on a 14/10 h light/dark cycle with controlled temperature and humidity. After 2 days of acclimation phase, Heme/PPIX (Frontier Scientific, USA) were administered intraperitonially (i.p. 30 mg/Kg). The recording continued for the following 7 days. The system consists of a standard GM-500 cage with a food hopper and a water bottle connected to load cells (2 mg precision) with 1 Hz rate data collection. Additionally, the cage contains a red house enrichment. Ambulatory activity was monitored at 1 Hz rate using an XY beam break array (1□cm spacing). Oxygen, carbon dioxide and water vapor were measured using a CGF unit (Sable Systems). This multiplexed system operated in pull□mode. Air flow was measured and controlled by the CGF (Sable Systems) with a set flow rate of 2□L/min. Oxygen consumption and carbon dioxide production were reported in milliliters per minute (mL/min). Energy expenditure was calculated using the Weir equation (Weir, 1949) and Respiratory Exchange Ratio (RER) was calculated as the ratio of VCO_2_/VO_2_. Raw data was processed using Macro Interpreter v2.41(Sable Systems). Similar set-up was used to assess the metabolic parameters of *Pcc* infected mice, except for the use of a 12/12 h light/dark cycle. Briefly, naïve *G6pc1*^*Alb*Δ/Δ^ and *G6pc1*^*fl/fl*^ mice were allowed to acclimatize within the cages for 3 days, and were taken out for infection. Mice were subsequently returned to the cages and monitored from days 6-9 of infection.

##### Luciferase Assays

HepG2 cells were seeded onto 6-well plates and transfected 24h thereafter (60%– 80% confluence) with full or truncated forms of pGL2B-G6PC (containing −1320/+60, −320/+60, −80/+60, from the rat *G6pc1* promoter, kind gift from Amandine Gautier-Stein; Université Claude Bernard Lyon, France)(Rajas et al., 2002) or the empty vector pGL2B alone using Lipofectamine LTX Reagent with PLUS Reagent (Invitrogen, Thermo Scientific), according to manufacturer’s instructions. A Renilla Luciferase-expressing vector (kind gift from Amandine Gautier-Stein; Université Claude Bernard Lyon, France)(Rajas et al., 2002) was co-transfected as transfection control. Forty-eight hours thereafter cells were treated with heme (40µM; 16h; 10% FBS 1%PS DMEM) and collected for luciferase activity assessment, determined using the Luciferase assay system Dual-Glo (Promega), according to manufacturer’s instructions. Luminescence was measured using a microplate reader (Glomax, Promega). Firefly Luciferase was normalized to Renilla and Luciferase activity is expressed as relative light units (RLU) using pGL2B activity as background, essentially as described (Rajas et al., 2002).

##### Primary Hepatocyte isolation, culture and heme exposure

Primary mouse hepatocytes were isolated, essentially as described (Dentin et al., 2004). Briefly, C57BL/6 mice were anesthetized (200µl/mouse, i.p., consisting of 30µl Imalgene 1000; MERIAL #915264 and 2µl Rompun 2%; Bayer #7427831 in ddH_2_O) and perfused with liver perfusion medium (25 mL; Gibco; 5 minutes) followed by freshly prepared collagenase buffer (42.5 mL HBSS; Gibco; 5mL 6.67mM CaCl_2_, 2.5 mL 1M Hepes, 12mg Collagenase, Sigma; 8 minutes). The digested liver was dissected to obtain crude liver single cell suspensions and hepatocytes were re-suspended in Percoll-based gradient (9.6 ml; 0.96 mL PBS 10X; 8,64 mL Percoll; GE), centrifuged (50*g*; 10 minutes). Hepatocytes were seeded on pre-coated 6 well plates (50 µg/mL rat collagen I, ThermoFisher in 20 mM acetic acid) at a density of 3×10^5^ cells per well in culture medium (William’s E Medium; 5% FBS; 1% Penicillin/Streptomycin). Culture medium was replaced after 4 hours and cells were left untouched for 20 hours before adding fresh medium containing heme (10 µM; Frontier Scientific, prepared in NaOH and neutralized with HCl) and/or Dexamethasone (Sigma Aldrich; 100 mM in 100% Ethanol, diluted in ddH2O). Cells were rinsed with PBS and harvested for RNA extraction and analysis.

##### Serology

Mice were sacrificed 7 days after *Plasmodium* infection and blood was collected for hemogram and serological analyses, performed by DNAtech (Portugal; http://www.dnatech.pt/web/).

##### Histology

Mice were sacrificed 7 days after *Plasmodium* infection, perfused *in toto* with ice-cold PBS and organs were harvested, fixed in 10% formalin, embedded in paraffin, sectioned (3μm) and stained with Hematoxylin & Eosin (H&E). Whole sections from formalin fixed and H&E-stained tissues were analyzed. Slides were analyzed with a DMLB2 microscope (Leica), and images were acquired with a DFC320 camera (Leica) and NanoZoomer-SQ Digital slide scanner (Hamamatsu Photonics). Histology was analyzed at the Instituto Gulbenkian de Ciência Histopathology Unit. Images were analyzed and constructed using the NDP.view2 (Hamamatsu Photonics) software.

##### Flow cytometry

Mice were sacrificed, perfused *in toto* (PBS; 10 mL) and spleens were collected in RMPI 1640, finely minced and passed through 70- and 40-μm nylon meshes. RBCs were lysed (5min, RT) in RBC lysis buffer (9 parts of 0,16M NH_4_Cl; + 1 part 0,1M Tris HCL pH 7,5). Cells were collected, washed (2x; ice cold PBS) and incubated with Fc Block (anti-CD16/CD32, clone 2.4G2, produced in house; 15 min.; 4°C) to prevent unspecific binding and with LIVE/DEAD™ Fixable Yellow Dead Cell Stain Kit (Invitrogen™), excluding dead cells from analysis. Cells were washed (ice cold PBS) and stained (20 min, 4°C) with fluorochrome-conjugated antibodies against: Ly6G (clone 1A8, PE-conjugated, BD Pharmingen™), TCRβ (clone H57-597; BV421-conjugated, Biolegend), CD3 (clone 17A2; BV711-conjugated, Biolegend), NK1.1 (clone PK136; A647-conjugated, produced in house), CD4 (clone GK1.5, Pe/Cy7-conjugated, Biolegend; or FITC-conjugated, produced in house), CD8 (clone 53-6.7; PercP/Cy5.5-conjugated, eBioscience™; or clone YTS169.4; PE-conjugated, produced in house), CD44 (clone IM7; eFluor 450-conjugated, eBioscience™), CD62L (clone Mel14; PercP/Cy5.5-conjugated, Biolegend), CD11b (clone M1/70, FITC-conjugated, BD Pharmingen™), CD19 (clone 6D5; BV711-conjugated, Biolegend), CD11c (clone N418; BV605-conjugated, Biolegend), F4/80 (clone BM8; A700-conjugated, Biolegend), CD163 (clone TNKUPJ; PercP-eFluor 710-conjugated, eBioscience™) and CD169 (clone 3D6.112; A647-Conjugated, Biolegend). Antibodies produced in house were purified from the respective hybridoma and, if applicable, labeled at the IGC’s Flow Cytometry and Antibodies Facility. Cells were analyzed on a LSR Fortessa X-20 (BD Biosciences). FACS data was analyzed with FlowJo V10.

For analysis of *P. falciparum* iRBC, *Pf*-3D7-GFP cultures were collected, washed (PBS) and stained with MitoTracker™ Deep Red FM (100nM in PBS; Invitrogen™, Thermo Fisher Scientific) for 15min. 37ºC. Cells were washed (PBS) again and stained with Hoechst 33342 (4mM in PBS; Thermo Fisher Scientific) for 15min. 37ºC, washed (PBS) and analyzed in a FACSAria™ Ilu Cell Sorter (BD Biosciences). FACS data was analyzed with FlowJo V10.

##### Cytokines

Mice were sacrificed 7 days after *Pcc* infection and plasma was collected from whole blood, for cytokine analyses, performed by Eve Technologies (Canada; https://www.evetechnologies.com/).

##### Western Blots

Proteins were extracted, electrophoresed and electro-transferred, essentially as previously described (Gozzelino et al., 2012; Ramos et al., 2019). Briefly, after euthanasia and perfusion (with ice-cold 1X PBS), livers were collected and snap frozen in liquid nitrogen. Tissue was lysed (2x SDS page sample buffer; 20% glycerol, 4% SDS, 100mM Tris pH6.8, 0.002% bromophenol blue, and 100mM dithiothreitol) and homogenized in a tissue lyser (Qiagen) with tungsten carbide beads (Qiagen). Supernatants were collected and total protein was quantified (λ_280nm_; Nanodrop 2000; ThermoFisher Scientific). Protein (100 μg) was resolved on a 12% SDS-PAGE and transferred to Polyvinylidene fluoride (PVDF) membranes. Membranes were blocked with 5% milk (in 1X T-TBS), washed with 1xT-TBS and incubated with primary antibodies overnight (4°C). Primary antibodies include anti-mouse G6PC1 (Rabbit, 1:5000)(Weis et al., 2017) and anti-GAPDH (Sicgen, Goat, 1:5000). Membranes were blocked (5% milk in 1X T-TBS) and were incubated with corresponding peroxidase-conjugated secondary antibodies (Goat: Donkey anti-Goat IgG (H+L) Secondary Antibody, HRP #PA1-28664; Rabbit: Goat anti-Rabbit IgG (H+L) Secondary Antibody, HRP #31460, Mouse: Peroxidase AffiniPure Donkey Anti-Mouse IgG (H+L) #715-035-150) for 1 hour at room temperature. Peroxidase activity was detected using SuperSignal™ West Pico PLUS Chemiluminescent Substrate (ThermoFisher Scientific). Blots were developed using Amersham Imager 680 (GE Healthcare), equipped with a Peltier cooled Fujifilm Super CCD. Densitometry analysis was performed with ImageJ (Rasband, W.S., ImageJ, U.S. NIH, Bethesda, Maryland, USA, https://imagej.nih.gov/ij/,1997-2014), using only images without saturated pixels.

##### RNA extraction and qRT-PCR

Mice were sacrificed, perfused *in toto* (10mL ice-cold PBS) and organs were harvested, snap frozen in liquid nitrogen and stored at -80ºC. Total RNA was extracted using tripleXtractor reagent (Grisp), chloroform (Merk) and with the NucleoSpin RNA kit (Machery-Nagel), according to manufacturer’s instructions. When indicated, whole blood was collected into heparinized tubes and RNA extracted using the NucleoSpin RNA Blood kit (Machery-Nagel), according to the manufacturer’s instructions. For mouse primary hepatocytes, cells were collected into tripleXtractor reagent (Grisp) and RNA was extracted using chlorophorm, isopropanol and ethanol, according to manufacturer’s instructions. cDNA was synthesized (Transcriptor First Strand cDNA Synthesis Kit, Roche or with Xpert cDNA Synthesis Kit, Grisp) for SYBR Green (iTaq Universal SYBR Green Supermix, Bio-Rad) based qPCR on a QuantStudio™ 7 Flex Real-Time PCR System (Applied Biosystems). Relative gene expression was calculated using the 2^−Δ^CT method (relative number) using Acidic ribosomal phosphoprotein P0 (*Arbp0*) as the housekeeping control gene for mouse gene or *Pcc 18S* for parasite genes. For list of PCR primers, see *key resource table*.

##### Bulk RNA sequencing and Analysis of liver samples

Livers were harvested from euthanized C57BL6 mice, 7 days after *Pcc* AS infection (*see Plasmodium infections*), 15 hours after heme administration (i.p., 30 mg/Kg BW) or after overnight fasting (controls), grinded and snap frozen in liquid nitrogen and stored at -80°C. RNA was extracted, cleaned (RNeasy MinElute Cleanup Kit, Qiagen) and assessed for quality using Agilent Bioanalyzer 2100 (Agilent Technologies) in combination with the RNA 6000 pico kit (Agilent Technologies). Full-length cDNAs and sequencing libraries were generated following the SMART-Seq2 protocol, as previously described (Picelli et al., 2014). After quality control (Agilent Technologies), library preparation including cDNA ‘tagmentation’, PCR-mediated adaptor addition and amplification of the adapted libraries was done following the Nextera library preparation protocol (Nextera XT DNA Library Preparation kit, Illumina), as previously described (Baym et al., 2015). Libraries were sequenced (NextSeq500 sequencing; Illumina) using 75 SE high throughput kit. Sequence information was extracted in FastQ format, using Illumina’s bcl2fastq v.2.19.1.403, producing around 29.10^6^ reads per sample. Library preparation and sequencing were optimized and performed at the Instituto Gulbenkian Ciência Genomics Unit.

The fastq reads were aligned against the mouse reference genome GRCm38 using the annotation GENCODE M25 to extract information about splice junctions (STAR; v.2.7.3a) (Dobin et al., 2013). FeatureCounts (v.2.0.1) was used to perform read summarization by assigning uniquely mapped reads to genomic features. Gene expression tables were imported into the R programming language and environment (v.3.6.3) to perform differential gene expression and functional enrichment analyses, as well as data visualization.

Differential gene expression was performed using the DESeq2 R package (v.1.31.11). Gene expression was modeled by *∼Condition* which included the following factor samples: *Pcc* infection (n=4), Heme administration (n=3) and Fasting (n=4). Genes not expressed or expressed less than 10 counts across the 11 samples were removed resulting in 18,136 genes for downstream differential gene expression analysis. We subsequently ran the function *DESeq* which estimates the size factors (by *estimateSizeFactors*, dispersion (by *estimateDispersions*) and fits a binomial GLM fitting for βi coefficient and Wald statistics (by *nbinomWaldTest*). Finally, the pairwise comparisons tested through contrasts with the function results, given the alpha of 0.05, were: Pcc *vs*. Fasting and Heme *vs*. Fasting. In addition, the log2 fold change for each pairwise comparison was shrunken with the function *lfcShrink* using the algorithm *ashr* (v.2.2-47)(Liao et al., 2014)). Differentially expressed genes are genes with an adjusted p-value<0.05 and an absolute log2 fold change>0. Normalized gene expression counts were obtained with the function *counts* using the option normalized = TRUE. Regularized log transformed gene expression counts were obtained with *rlog*, using the option blind = TRUE, for Principal Component Analysis of overall sample expression profiles and for hierarchical clustering of differentially expressed genes.

The Ensembl gene ids were converted into gene symbols from Ensembl (v.101 - Aug 2020-https://aug2020.archive.ensembl.org) by using the mouse reference (GRCm38.p6) database with biomaRt R package (v.2.46.2). All scatter plots, including volcano plots, were done with the ggplot2 R package (v.3.3.2). Venn diagrams were made with VennDiagram (v.1.6.20). Heatmaps were made with ComplexHeatmap (v.2.2.0), using the default options for distance and clustering method estimation.

Functional enrichment analysis was performed with the gprofiler2 R package (v.0.2.0). The function *gost* was applied in order to perform enrichment based on the list of up- or down-regulated genes (genes with an adjusted p-value<0.05 and a log2 fold-change>0 or <0), between each pairwise comparison (independently), against the annotated genes (domain_scope = “annotated”) of the organism Mus musculus (organism = “mmusculus”). The gene lists were ordered by increasing adjusted p-value (ordered_query = TRUE) in order to generate a GSEA (Gene Set Enrichment Analysis) style *p*-values. In addition, only statistically significant (user_threshold=0.05) enriched functions are returned (significant=TRUE) after multiple testing corrections with the default method g:SCS (correction_method = “g_SCS”). gprofiler2 was run against all the default functional databases for mouse which include: Gene Ontology (GO:MF, GO:BP, GO:CC), KEGG (KEGG), Reactome (REAC), TRANSFAC (TF), miRTarBase (MIRNA), Human phenotype ontology (HP), WikiPathways (WP), CORUM (CORUM). gprofiler2 was with the archived version of the gprofiler2 server to ensure reproducibility - Ensembl 102, Ensembl Genomes 49 (database built on 2020-12-15): https://biit.cs.ut.ee/gprofiler_archive3/e102_eg49_p15/gost.

##### Transcription factor binding motif prediction

The -320 to +20 bp promoter region sequence of the mouse *G6pc1* was entered into the JASPAR database (http://jaspar.genereg.net), a non-redundant set of transcription factor binding profiles derived from published datasets of transcription factors binding sites (Sandelin et al., 2004). The transcription factors SP1, SP3, KLF3, KLF5, KLF7, ZF5, E2F-1, E2F-3, E2F-4, KROX/EGR1, CTCF or WT1 and AhR were assessed for predicted binding sites. Candidate binding sites were identified with predicted motifs using a relative profile score threshold of 80% (*i*.*e*., the default JASPAR setting). Motif scores are JASPAR relative scores, which is defined as 1 for the maximum-likelihood sequence.

##### Single-cell RNA sequencing

Blood was collected from *G6pC1*^*fl/fl*^ or *G6pc1*^*Alb*Δ/Δ^ mice at the peak of infection (*i*.*e*. day 7) with a transgenic *Pcc* AS expressing GFP. Samples were diluted 1:7.5 in 1xPBS and iRBCs were sorted according FSC, SSC and GFP on a BD FACS Aria Ilu. Cells were washed twice with 1x PBS, counted, and adjusted to a concentration of 1200 cells/µL.

##### Single-cell gene expression library preparation and sequencing

Cell suspensions were processed with Chromium™ Single Cell Controller to generate barcoded single-cell gel bead emulsions (GEMs) following the Chromium Next GEM Single Cell 3□ protocol. The single-cell cDNA libraries were obtained after the GEM-RT (reverse-transcriptase) clean-up and cDNA amplification on a Bio-Rad C1000 Touch™ Thermal Cycler with the following PCR thermal program: 1) 98 ºC for 3 min, 2) 98 ºC for 15 sec, 3) 63 ºC for 20 sec, 4) 72 ºC for 1 min, (Steps 2 to 4 cycled 11 x), 5) 72 ºC for 1 min and 6) 4 ºC Hold. The cDNA profiles were checked on a Fragment Analyzer System (Agilent Technologies) according to the HS NGS Fragment kit (Agilent Technologies) manual. The scRNA-seq libraries were generated from 10µl (25%) of the single-cell cDNA sample and indexed with the Chromium i7 Multiplex Kit (10x Genomics) on a Bio-Rad C1000 Touch™ Thermal Cycler as follows: 1) 98 ºC for 45 sec, 2) 98 ºC for 20 sec, 3) 54 ºC for 30 sec, 4) 72 ºC for 20 sec, (Steps 2 to 4 cycled 15 x), 5) 72 ºC for 1 min and 6) 4 ºC Hold. The final 3’ Gene Expression libraries were verified and quantified on a TapeStation 4200 using the D1000 ScreenTape kit (Agilent Technologies). Samples were sequenced on a NextSeq 500 (Illumina) using the NextSeq 500/550 High Output Kit v2.5 (150 Cycles) with the following sequencing run settings: Read1: 28cycles, i7 index: 8 cycles, i5 index 0 cycles, Read2: 130 cycles, aiming a read depth of _∼_25,000 reads/cell and.

##### Single-cell RNA-seq raw data analysis

Illumina sequencer base call files (BCL) were processed using Cell Ranger mkfastq pipeline generating FASTQ files. A custom genome reference was created for *Plasmodium chabaudi* following the steps described in https://support.10xgenomics.com/single-cell-gene-expression/software/pipelines/latest/using/tutorial_mr. Briefly a GTF file was created from the *Plasmodium chabaudi* GFF file and genome deposited at ftp://ftp.sanger.ac.uk/pub/genedb/releases/latest/Pchabaudi (retrieved on 2021, February 21) using the cellranger mkgtf command with the option to filter coding proteins. After, the reference genome was created by merging the GTF file with the genome of *P. chabaudi* in FASTA format using the cellranger mkref command. The two expression libraries were then analyzed with the Cell Ranger count pipeline v4.0.0 using the created *P. chabaudi* reference genome. All further analyses were performed using R (v. 3.6.3; R Core Team 2020). Samples were analyzed with the Seurat R package (v.4.0.0) (Butler et al., 2018; Hao et al., 2020; Satija et al., 2015) following the procedures of https://satijalab.org/seurat/articles/pbmc3k_tutorial.html. Samples were processed independently before integration as described https://satijalab.org/seurat/articles/integration_introduction.html. First, genes not expressed at least in 3 cells were removed as well as cells with a number of expressed genes higher and equal than 750 and a total number of reads higher or equal than 1500. These filters resulted in 4554 genes and 15537 cells for sample *G6pc1*^*Alb*Δ/Δ^ and 4545 genes and 5176 cells in the case of sample *G6pc*^*lox/lox*^. Both objects were normalized, variable features were determined, with the threshold of 750 most variable features and integrated features were selected. Integration was performed by finding shared anchors across the samples using the integrated variable features found before. The integrated Seurat object was re-scaled, PCA and UMAP were determined by using the first 12 principal components (PCs). Neighbors were found based on the first 12 PCs and PCA as the reduction method. Clustering was performed using the first 12 PCs and a resolution of 0.39 (based on the resolution used for the independent analysis of the sample of iRBCs of *G6pc1*^*Alb*Δ/Δ^), which yielded 9 clusters. The clusters were ascribed to different developmental stages of *Pcc*, according to the 25 highest conserved transcripts and by analogy to *P. falciparum* and *P. berghei* orthologues (Malaria Cell Atlas; https://www.sanger.ac.uk/tool/mca/). Differential expressed genes were identified using the Wilcoxon Rank Sum test and expression was analyzed across *Pcc* of iRBCs of *G6pc1*^*Alb*Δ/Δ^ *vs. G6pC1*^*fl/fl*^ for each cluster. It was not possible to test differential gene expression for cluster 8 as this cluster is almost exclusive of *Pcc* of iRBCs of *G6pc1*^*Alb*Δ*/*Δ^. Functional enrichment analysis between the differentially expressed up- and down-regulated genes obtained from each pairwise comparison between each cluster across the two samples was performed by the use of gprofiler2 R package (v.0.2.0) (Kolberg et al., 2020)(an interface to the g:Profiler web browser tool using the archived version of the gprofiler2 server - Ensembl 102, Ensembl Genomes 49 (database built on 2020-12-15): https://biit.cs.ut.ee/gprofiler_archive3/e102_eg49_p15/gost. Enrichment was performed based on the list of up- or down-regulated genes, between each pairwise comparison (independently), against the annotated genes of the organism *Plasmodium chabaudi available on the Gene Ontology database (GO:MF, GO:BP, GO:CC)*. The gene lists were ordered by increasing adjusted p-value to generate a GSEA (Gene Set Enrichment Analysis) style *p*-values, following the same procedure described for Bulk RNA sequencing and analysis of liver samples. Functional enrichment was tested for all clusters, except for cluster 8, as it lacks differential gene expression data. Genes were ranked up according to avg_log2FC > 0 or down avg_log2FC < 0. The ggplot2 R package was used for visualization (v.3.3.2) and the ComplexHeatmap package (v.2.2.0) for heatmaps.

For RNA-velocity a .loom file was created with the spliced and unspliced count matrices for each sample using the velocyto CLI tool (v.0.17.17)(La Manno et al., 2018) with the run10x command providing the previous GTF file of Pcc and the output of the Cell Ranger pipeline with the bam files and features matrix data. GNU parallel was used to run in parallel both samples (v.20161222). RNA velocity was estimated with the python package scvelo (v.0.2.2) using the stochastic model, and the velocities obtained were projected in the UMAP embeddings obtained previously with the Seurat software. Estimating RNA velocity from the spliced and unspliced matrices uses the ratio of spliced and unspliced RNA transcripts to determine the cell trajectories. Other packages used were os, random, anndata (v.0.7.5), pandas (v.1.1.3), numpy (v.1.19.2) and matplotlib (v.3.3.2) using python (v.3.8.3). This analysis is based on the tutorial: https://github.com/basilkhuder/Seurat-to-RNA-Velocity.

Similarly to the above-described analysis of *P. chabaudi*, the reads of the murine RBCs were also analyzed to verify that similar amounts of cells were used for sequencing. In this case, the first 4 PCs were used for the computation of the UMAP. Downstream analysis of clusters was not performed.

## Data and Software availability

Raw data files for the bulk and single cell RNA sequencing were deposited to BioStudies ArrayExpress database (https://www.ebi.ac.uk/arrayexpress/) with accession numbers E-MTAB-10935 (for liver bulk RNA) and E-MTAB-10939 (for single cell).

## Statistical analysis

Statistically significant differences between two experimental groups were assessed using a two-tailed unpaired Mann-Whitney U test or t-test. Comparisons between more than two groups were carried out by two-way ANOVA with Tukey’s or Sidak’s multiple-comparisons test, or Kruskal-Wallis test with Dunn’s multiple-comparisons test. Survival was represented by Kaplan–Meier plots and difference between the groups was assessed using the log-rank test. All statistical analyses were performed using GraphPad Prism 8 software. Differences were considered significant at a p-value <0.05. *ns*: Not significant, p>0.05; *: p<0.05, **: p<0.01; ***: p<0.001; ****: p<0.0001.

