## Supplementary figures and images for "A HYPOMETABOLIC DEFENSE STRATEGY AGAINST *PLASMODIUM* INFECTION"

### Figure S1

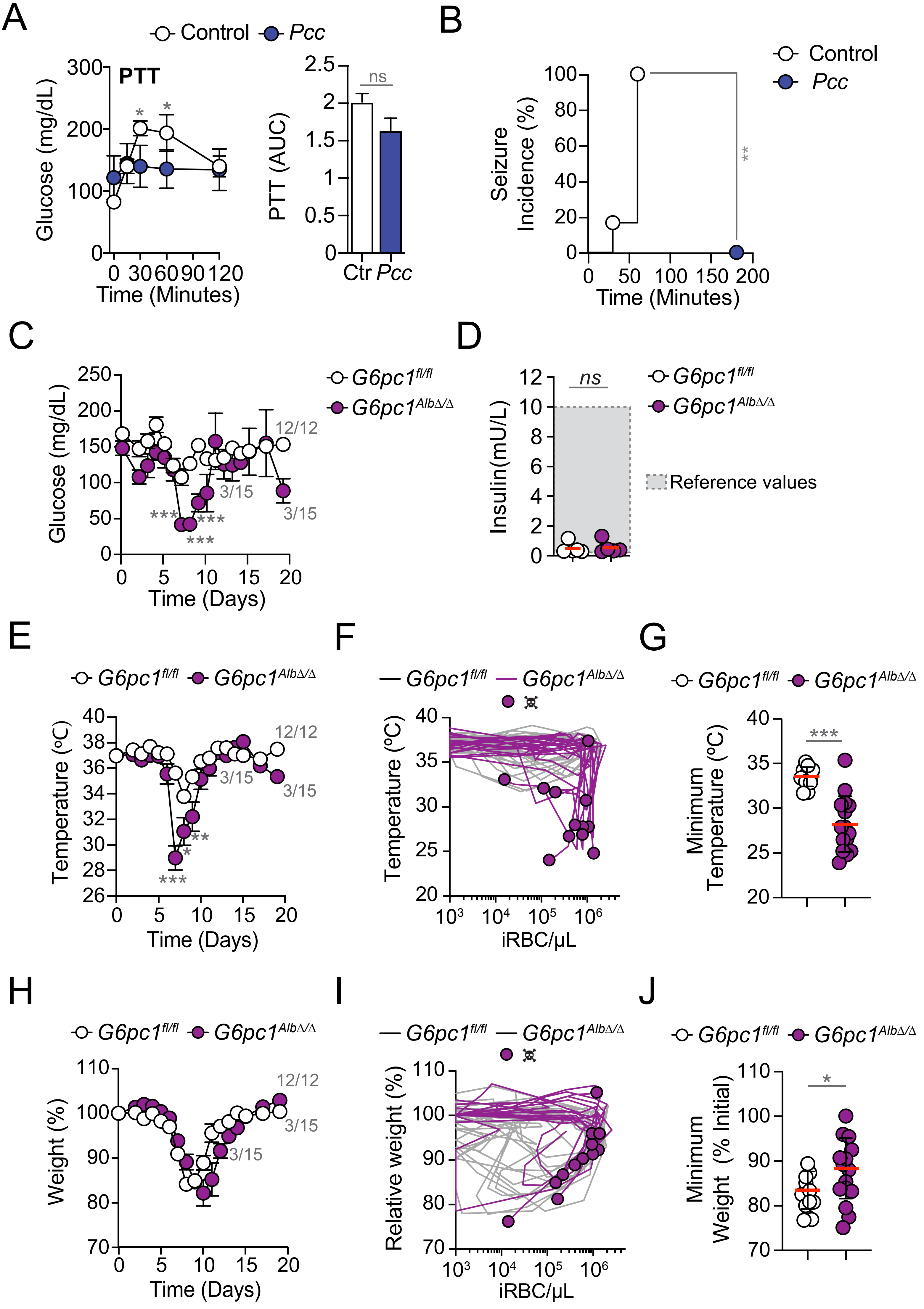

### Figure S2

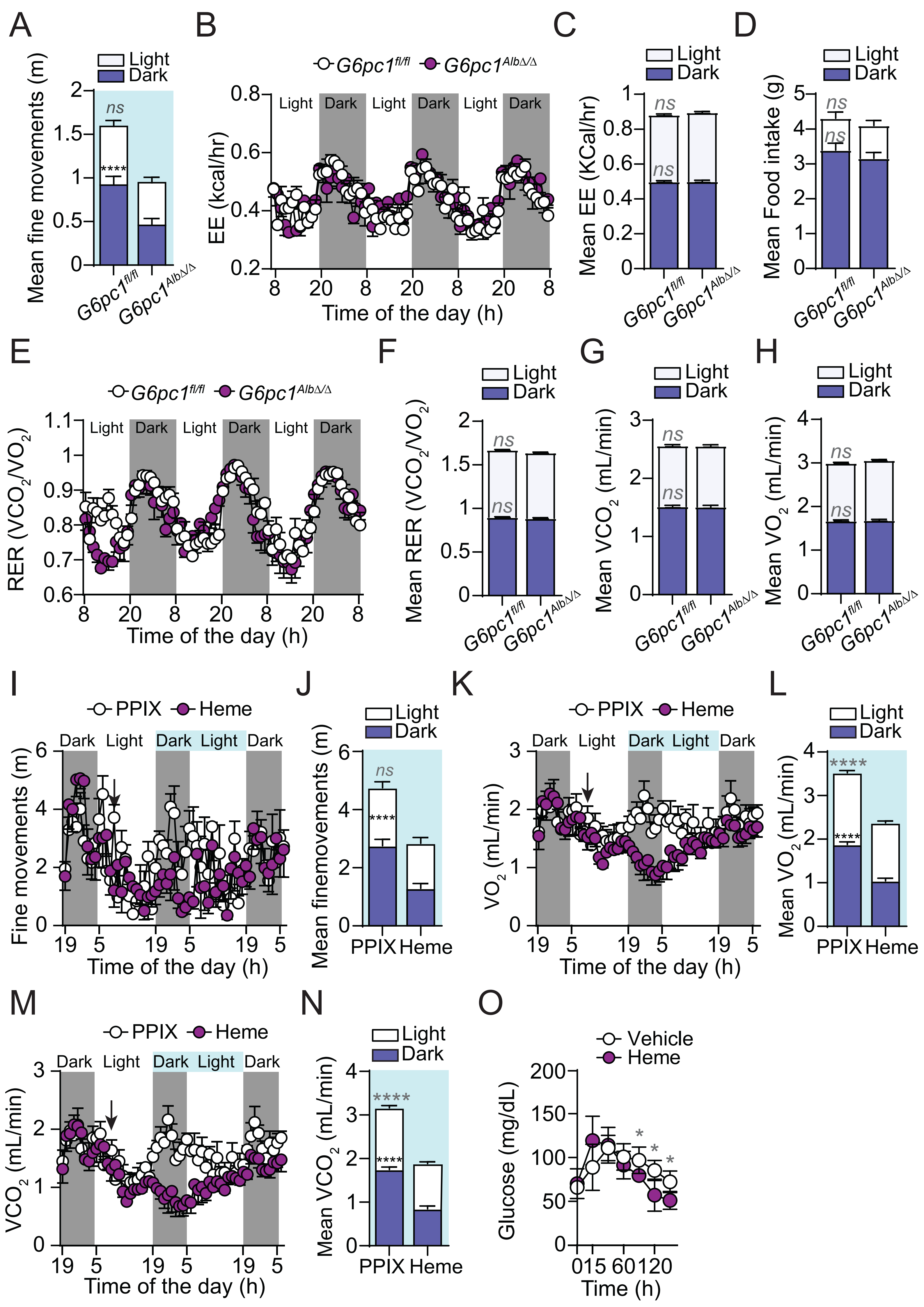

### Figure S5

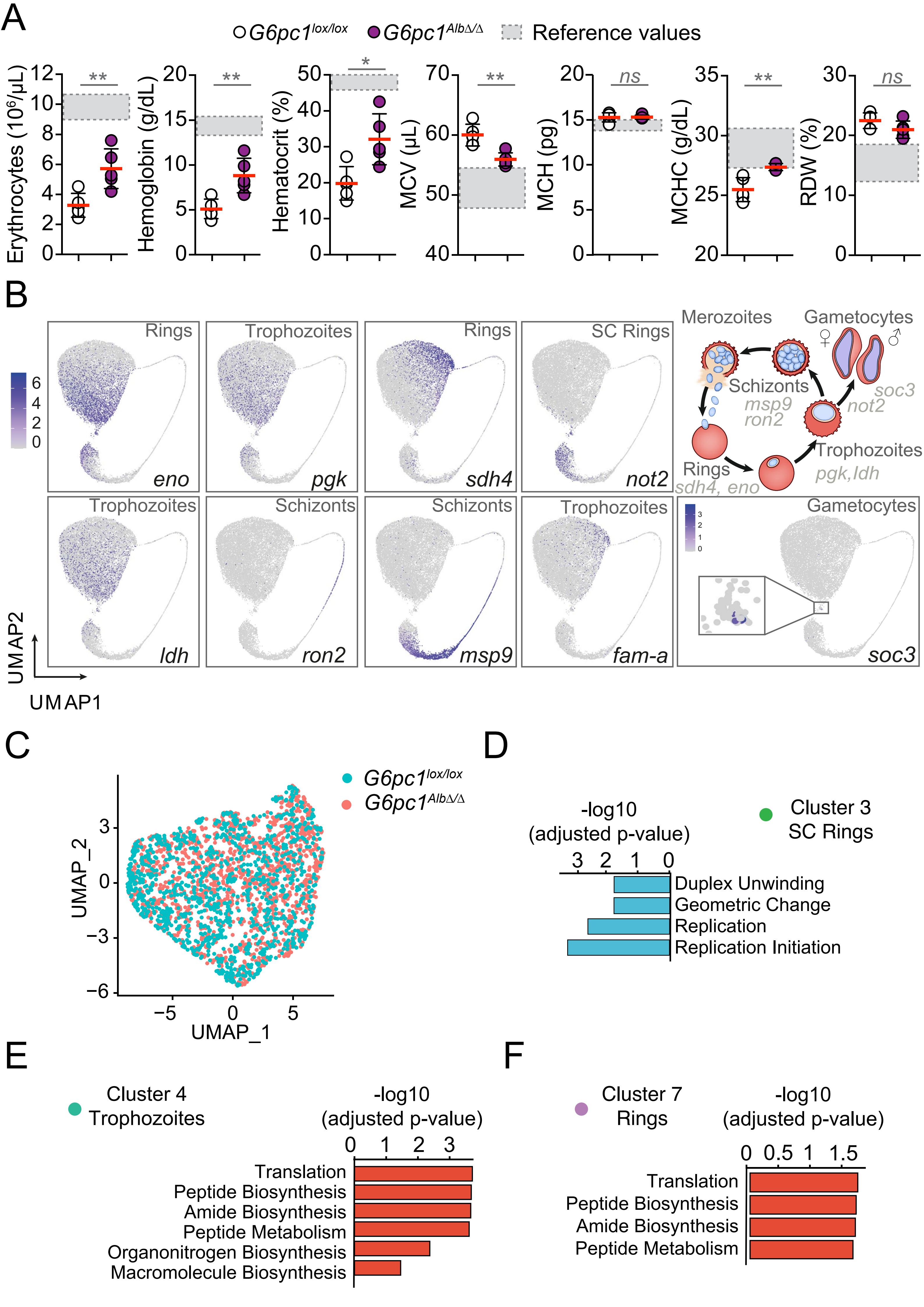

### Figure S6

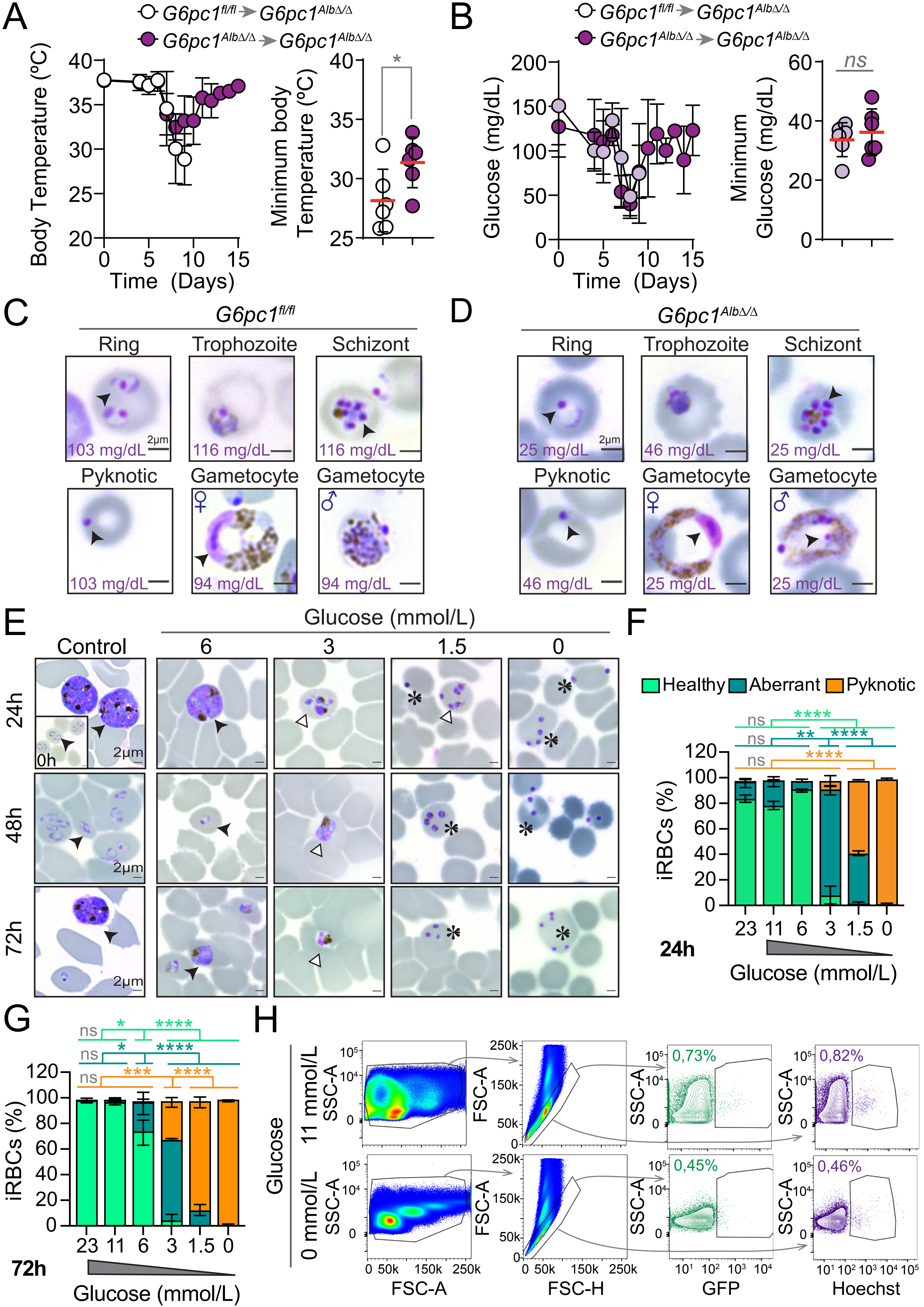

### Figure S7

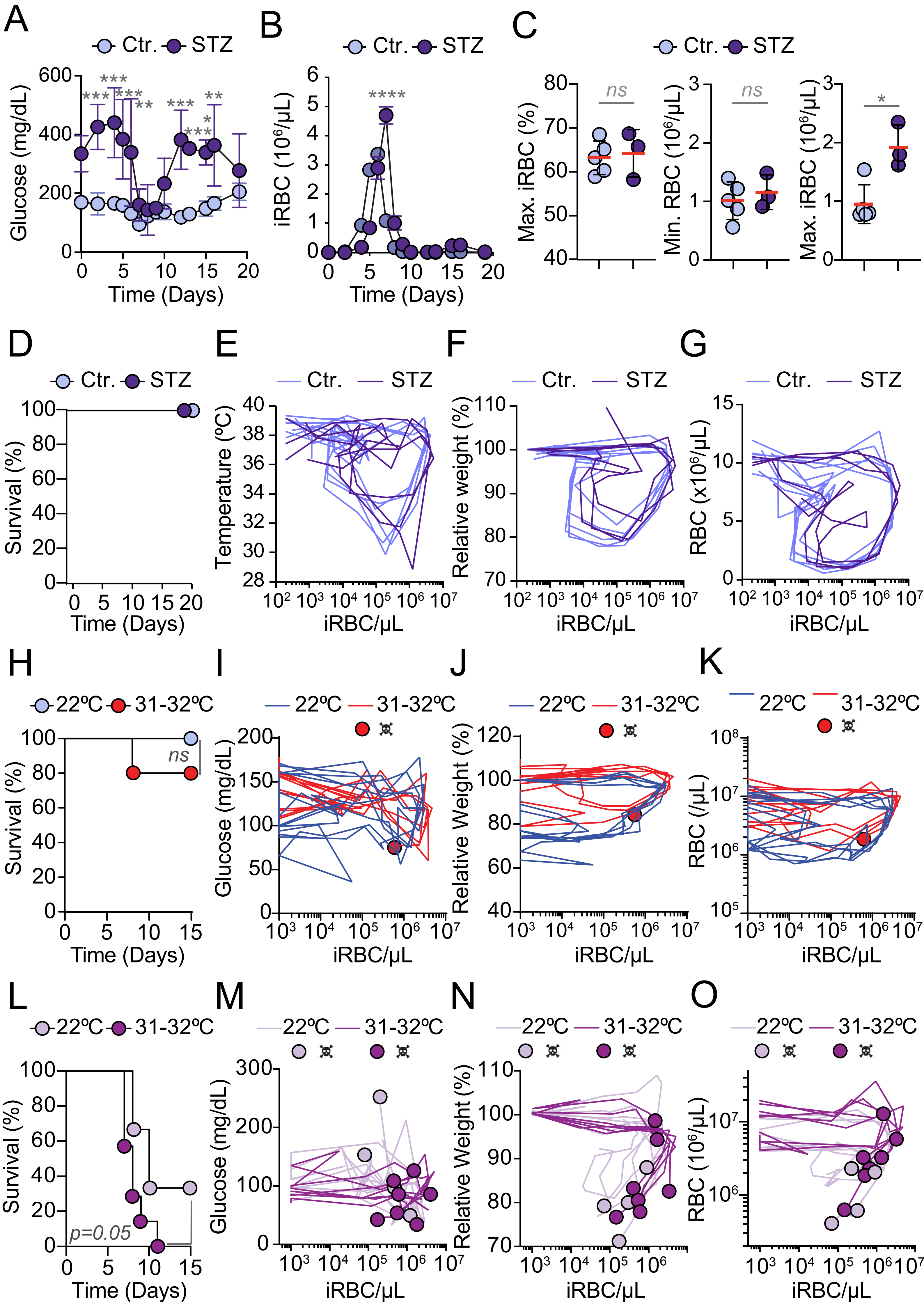
