## Supplementary material for "A HYPOMETABOLIC DEFENSE STRATEGY AGAINST *PLASMODIUM* INFECTION": Figure S3

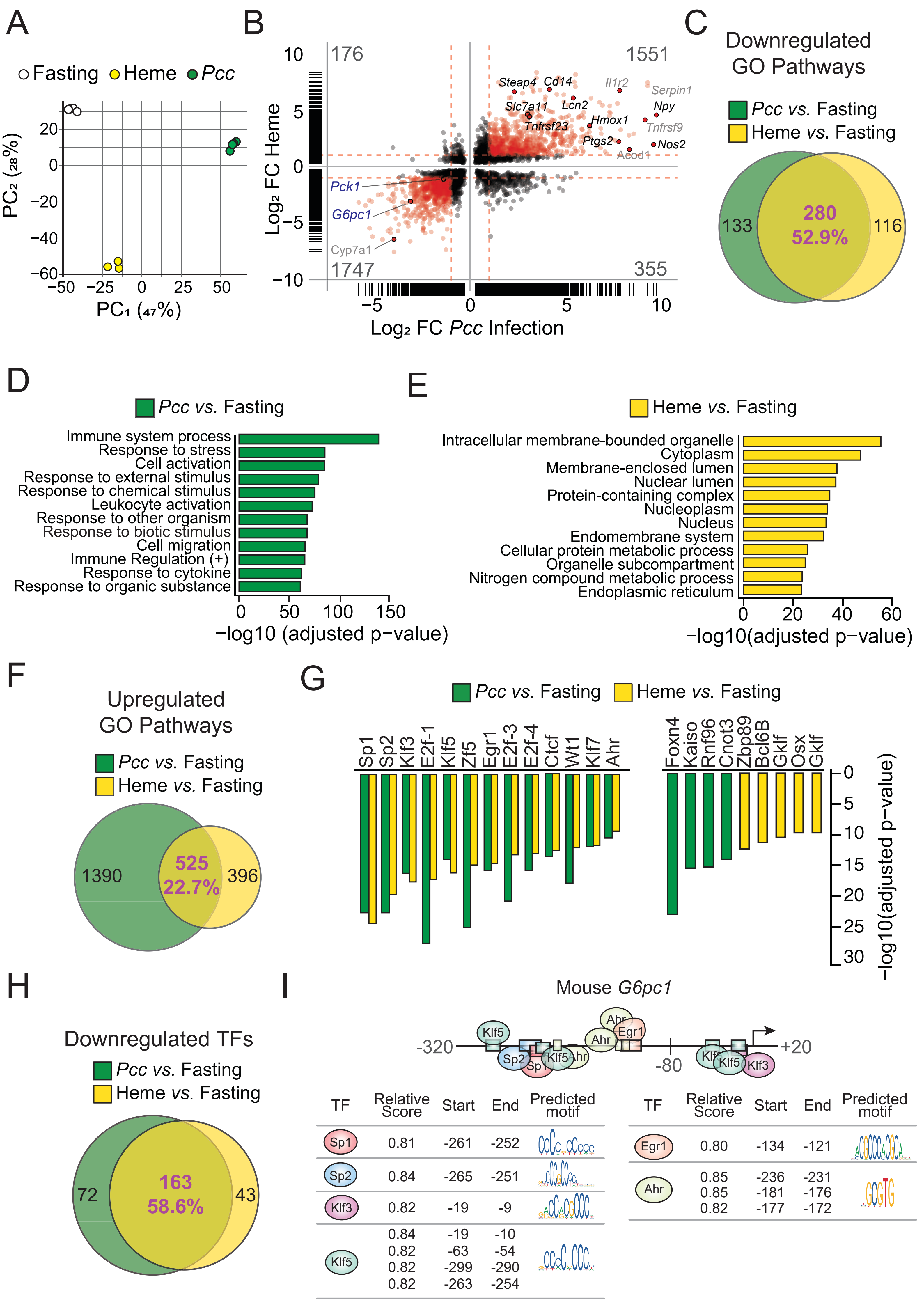


| TF | Relative Score | Start | End | Predicted motif |
|---|---|---|---|---|
| Sp1 | 0.81 | -261 | -252 | |
| Sp2 | 0.84 | -265 | -251 | |
| Klf3 | 0.82 | -19 | -9 | |
| Klf5 | 0.84<br>0.82<br>0.82<br>0.82 | -19<br>-63<br>-299<br>-263 | -10<br>-54<br>-290<br>-254 | |

| TF | Relative Score | Start | End | Predicted motif |
|---|---|---|---|---|
| Egr1 | 0.80 | -134 | -121 | |
| Ahr | 0.85<br>0.85<br>0.82 | -236<br>-181<br>-177 | -231<br>-176<br>-172 | |
