## Supplementary material for "A HYPOMETABOLIC DEFENSE STRATEGY AGAINST *PLASMODIUM* INFECTION": Figure S4

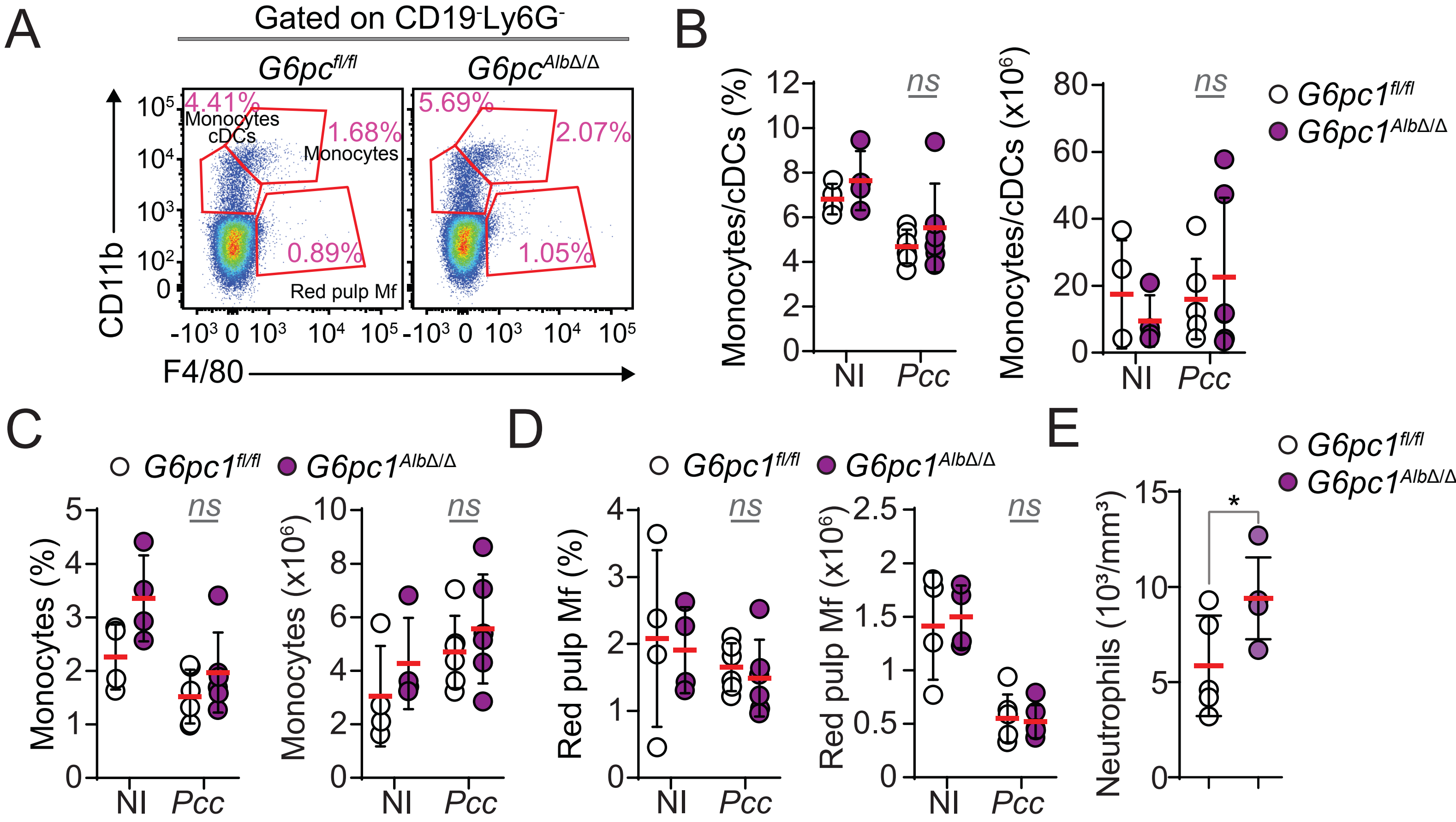


F

| | Naive | | Pcc | |
|---|---|---|---|---|
| | *G6pc1fl/fl* | *G6pc1AlbΔ/Δ* | *G6pc1fl/fl* | *G6pc1AlbΔ/Δ* |
| Eotaxin | 2571.7±632.4 | 3162.2±959.9 | 938.4±213.2 | 1508.9±309.7* |
| IL1α | 109.7±103.9 | 160.9±191.9 | 95.3±79.3 | 55.1±50.8 |
| IL1β | 33.3±54.6 | 4.2±5.1 | 2.5±3.0 | 2.5±1.7 |
| IL2 | 3.4±2.4 | 7.2±3.0 | 4.0±2.8 | 8.5±6.7 |
| IL3 | 0±0 | 1.5±1.3 | 1.5±2.3 | 1.7±1.1 |
| IL4 | 0.5±0.2 | 3.5±1.3 | 2.1±0.9 | 3.2±4.0 |
| IL5 | 13.5±2.2 | 27.6±18.3 | 9.3±6.6 | 5.8±5.7 |
| IL6 | 22.0±17.3 | 12.0±5.2 | 77.7±58.3 | 35.6±41.4 |
| IL7 | 6.4±4.6 | 7.2±4.6 | 9.2±6.2 | 12.8±10.6 |
| IL9 | 45.9±0 | 59.4±85.8 | 164.7±85.6 | 244.1±227.6 |
| IL10 | 12.0±3.9 | 22.1±12.7 | 964.8±968.3 | 722.2±627.3 |
| IL12p40 | 23.6±32.1 | 26.4±7.9 | 2.5±6.2 | 5.3±6.9 |
| IL12p70 | 20.9±5.2 | 82.7±112.1 | 8.8±11.5 | 4.0±5.4 |
| IL13 | 60.3±37.1 | 70.0±13.3 | 36.0±14.6 | 38.5±8.7 |
| IL15 | 117.4±98.0 | 91.5±9.9 | 259.7±153.2 | 143.7±130.4 |
| IL17 | 3.1±0.8 | 5.8±1.5* | 0.5±1.4 | 0.2±0.4 |
| Cxcl10 | 74.0±24.1 | 103.5±23.75 | 758.8±305.2 | 547.6±319.3 |
| KC | 70.4±94.2 | 138.3±155.3 | 82.6±67.0 | 43.4±46.2 |
| Lif | 0.9±1.3 | 1.8±0 | 7.2±5.4 | 3.8±3.6 |
| Cxcl5 | 279.9±51.9 | 916.6±854.0* | 124.6±136.1 | 84.9±92.7 |
| Ccl2 | 9.3±11.5 | 916.6±854.0 | 8.7±21.5 | 4.4±7.9 |
| Cxcl9 | 24.8±6.3 | 35.0±11.9 | 592.2±263.4 | 634.4±208.2 |
| Ccl3 | 61.6±45.0 | 89.7±49.2 | 97.8±62.1 | 43.1±31.0 |
| Ccl4 | 59.1±46.0 | 78.4±36.8 | 524.6±453.7 | 255.8±202.7 |
| Cxcl2 | 130.1±151.0 | 81.9±60.8 | 87.1±99.6 | 52.5±66.7 |
| Ccl5 | 68.3±44.2 | 116.0±66.0 | 173.1±126.3 | 105.8±41.3 |
| Cxcl21 | 2040.0±160.9 | 1958.6±1575.6 | 4286.0±3158.3 | 5034.4±2992.1 |
| Fractalkine | 275.0±23.6 | 309.2±187.8 | 383.0±108.1 | 403.3±156.2 |
| IFNβ | 49.5±26.8 | 73.4±17.6 | 160.1±80.9 | 119.5±97.7 |
| IL-11 | 3.2±7.1 | 0±0 | 72.7±60.2 | 55.5±70.3 |
| IL-16 | 1190.6±392.0 | 1526.9±706.5 | 3110.5±1697.4 | 3294.5±1983.2 |
| IL-20 | 193.5±188.6 | 289.5±71.4 | 128.9±181.9 | 212.0±179.4 |
| Ccl22 | 207.5±56.1 | 185.3±95.7 | 90.7±62.3 | 105.9±55.1 |
| Ccl12 | 24.1±17.9 | 39.3±26.7 | 92.4±64.7 | 89.2±58.4 |
| Ccl20 | 108.4±147.7 | 101.2±82.7 | 12.2±10.6 | 14.0±14.8 |
| Ccl19 | 90.2±62.1 | 341.6±189.6 | 57.6±47.7 | 33.7±43.1 |
| Ccl17 | 47.4±36.5 | 50.4±34.24 | 29.7±12.5 | 33.34±23.0 |
| TIMP1 | 618.7±349.3 | 1279.5±1029.6 | 3293.2±1263.1 | 3967.3±2775.6 |
